# Identification of a cardiac glycoside exhibiting favorable brain bioavailability and potency for reducing levels of the cellular prion protein

**DOI:** 10.1101/2022.08.22.504810

**Authors:** Shehab Eid, Thomas Zerbes, Declan Williams, Xinzhu Wang, Chris Sackmann, Sammy Meier, Nickolai O Dulin, Pavel Nagorny, Gerold Schmitt-Ulms

## Abstract

Several strands of investigation have established that a reduction in the levels of the cellular prion protein (PrP^C^) is a promising avenue for the treatment of prion diseases. We recently described an indirect approach for reducing PrP^C^ levels that targets Na,K-ATPases (NKAs) with cardiac glycosides (CGs), causing cells to respond with the degradation of these pumps and nearby molecules, including PrP^C^. Because the therapeutic window of widely used CGs is narrow and their brain bioavailability is low, we set out to identify a CG with improved pharmacological properties for this indication. Starting with the CG known as oleandrin, we combined *in silico* modeling of CG binding poses within human NKA folds, CG structure-activity relationship (SAR) data, and predicted blood-brain barrier (BBB) penetrance scores to identify CG derivatives with improved characteristics. Focusing on C4’-dehydro-oleandrin as a chemically accessible shortlisted CG derivative, we show that it reaches four times higher levels in the brain than in the heart one day after subcutaneous administration, exhibits promising pharmacological properties, and suppresses steady-state PrP^C^ levels by 84% in immortalized human cells that have been differentiated to acquire neural or astrocytic characteristics. Finally, we validate that the mechanism of action of this approach for reducing cell surface PrP^C^ levels requires C4’-dehydro-oleandrin to engage with its cognate binding pocket within the NKA α subunit. The improved brain bioavailability of C4’-dehydro-oleandrin, combined with its relatively low toxicity, make this compound an attractive lead for brain CG indications and recommends its further exploration for the treatment of prion diseases.

**AUTHOR SUMMARY:** Prion diseases are fatal neurodegenerative diseases for which there is no effective treatment. An abundance of data indicates that reducing the levels of a specific protein, termed the cellular prion protein (PrP^C^), would not only be safe but would delay disease onset and extend prion disease survival. This project builds on our recent discovery that PrP^C^ binds to NKAs, specific cellular transport proteins that use energy to electrify cellular membranes by pumping charged potassium and sodium metals in and out of cells. We showed that targeting NKAs with their natural inhibitors, cardiac glycosides (CGs), causes brain cells to internalize and degrade NKAs, and that PrP^C^, on account of residing next to NKAs, gets co-degraded. Natural CGs act primarily on the heart. Here, we used computational modeling to identify a CG, termed KDC203, that is predicted to have favorable characteristics for brain applications. We show that KDC203 reduces PrP^C^ levels by 84% in immortalized human brain-like cells grown in the dish. Moreover, we show that KDC203 exhibits relatively low toxicity, predominantly targets the brain when subcutaneously injected into mice, and has other promising pharmacological characteristics that recommend it for further evaluation for the treatment of prion diseases.

## INTRODUCTION

The cellular prion protein (PrP^C^) is widely understood to undergo conformational conversions that underlie a group of neurodegenerative diseases, known as prion diseases [1]. Several lines of evidence also point toward PrP^C^ as a prominent receptor for the cellular docking of oligomeric Aβ peptides that mediate toxicity in Alzheimer’s disease [2].

Experimental ablation of the prion gene does not appear to confer severe phenotypes in mice [3] or cattle [4], a finding that has since been corroborated when a goat with a natural bi-allelic nonsense mutation in the prion gene was identified [5]. Consistent with these observations, human 23andMe data indicate that individuals with only one functional *PRNP* allele can reach advanced age in good health [6].

Critically, mice lacking the *Prnp* gene are refractory to prion disease, and *Prnp* heterozygosity approximately doubles the prion disease survival time [7]. Moreover, when PrP^C^ levels were suppressed in prion-infected mice after early spongiform degeneration or cognitive and neurophysiological prion disease symptoms were beginning to manifest, a partial rescue of these phenotypes could be observed [8, 9].

Consequently, reducing steady-state levels of PrP^C^ may be safe and may have merit for the treatment of prion diseases and Alzheimer’s disease.

To date, attempts to identify PrP^C^ lowering drugs through screens of compound libraries have largely failed, with some of the best lead compounds either requiring relatively high concentrations to exert their effect or lacking favorable ADME characteristics [10, 11]. Recent results from a study that targeted the stability of PrP^C^ transcripts by treating prion-infected mice with antisense oligonucleotides (ASOs) provided elegant proof-of-concept validation of the premise that lowering steady-state PrP^C^ levels can extend survival of prion-infected mice in a dose-dependent manner [12]. Robust data for the use of ASOs to address human brain disease exist in the form of advances in the treatment of spinal muscular atrophy [13]. Perhaps more relevant, the treatment of cynomolgus macaques with ASOs designed to induce the destruction of transcripts coding for the tau protein led to approximately 50% reductions in tau protein levels in several brain structures [14].

Challenges associated with adapting this approach for the treatment of human prion diseases relate to the need to inject ASOs periodically through the intrathecal route because mRNA levels are shown to recover two months following bolus injections [15], and limitations in the delivery of ASOs to deep brain structures, a caveat that may be exacerbated in human adults due to their relatively large brain sizes [16]. Thus, to date, no treatment has reached the clinic that can effectively reduce human brain PrP^C^ levels by targeting the expression or stability of this protein directly.

We recently discovered that PrP^C^ is surrounded by Na,K-ATPases (NKAs) in the brain [17], which led us to propose an indirect targeting approach [18]. We showed that when cardiac glycosides (CGs), a class of compounds also known as cardiotonic steroids, bind to NKAs, it causes their internalization and degradation. Rather than this representing a selective internalization of only the ligand-receptor complex, we observed that CG exposure of human neurons and astrocytes causes other NKA-associated molecules, including PrP^C^, to also be removed from the cell surface and degraded. A replacement of key amino acid residues in the CG-docking site of ATP1A1 in the human ReN VM cell model prevented the CG-dependent PrP^C^ degradation, indicating that the latter is dependent on CG ligand-ATP1A1 receptor engagement in this paradigm, as opposed to some other non-specific effect of CGs on the cell. Finally, we learned that the degradation of PrP^C^ under these circumstances relies on the endo-lysosomal system and specifically involves the cysteine protease cathepsin B [18].

CGs have a long history of use in the clinic for the treatment of heart diseases [19–21]. More recently, compounds from this class have been considered for other uses, including the treatment of cancers [22–24]. In addition, the notion to widen the use of CGs to the treatment of stroke and neurodegenerative diseases has had some traction [25]. To date, brain indications of CGs are limited by the relative narrow therapeutic window and poor blood-brain barrier (BBB) penetration or brain retention of members of this compound class [26, 27]. Arguably most attention in this regard has been paid to a CG known as oleandrin, which can be derived from the ornamental shrub *Nerium oleander* [28, 29] and has shown to accumulate to relatively high levels in brain tissue [30, 31]. However, oleandrin has recently been shown to exhibit inadvertent cardiotoxicity that exceeded the toxicity of other CGs and may limit its use [32].

We therefore set out to identify a CG with more favorable characteristics for the objective to lower steady-state PrP^C^ levels in the brain. To this end, we initially modeled the predicted binding of oleandrin to human NKAs expressed in the brain. Next, we considered available structure-activity relationship (SAR) data to narrow the possible chemical space for modifying oleandrin to those changes that can be attained with good yields. The *in silico* evaluation of a combinatorial CG library through a virtual screen revealed a small number of derivatives with promising BBB penetrance and docking scores. Subsequent work focused on C4’-dehydro-oleandrin, hereafter termed KDC203. We will show that KDC203 has similar potency as oleandrin in cell-based assays yet exhibits improved brain penetrance. In fact, 24 hours after subcutaneous injection brain levels of KDC203 were higher than its levels in heart, kidney, or liver. Moreover, we document that KDC203 is a lesser substrate than oleandrin of the human multi-drug transporter MDR1, known to be responsible for the rapid extrusion of other CGs, and is projected to reach an unbound concentration exceeding 10 nM in the brain. Finally, we show that exposure of cells of human neuronal and astrocytic lineage to low nanomolar KDC203 levels causes a profound reduction in the steady-state PrP^C^ levels through a mechanism of action that requires direct engagement of KDC203 with its NKA target.

## RESULTS

### Study design

A vast majority of commercially available CGs exhibit poor brain bioavailability, either because they are too hydrophilic or are actively extruded from the brain [33–35]. To our knowledge, no large libraries of less investigated CGs are commercially available. The *de novo* synthesis of CGs is challenging and requires dozens of steps [22, 36–39]. These realities dissuaded us from pursuing a screen-based approach for the identification of a CG that exhibits favorable characteristics for brain indications. Instead, we decided to build on an existing CG, oleandrin, that has shown promising brain bioavailability and employ a study design that pairs rational design with chemical derivatization to improve its properties for our indication. In this approach, a small number of compounds are shortlisted on the basis that they are predicted to have high blood-brain-barrier penetrance *and* are chemically accessible with high purity and yield through a small number of derivatization steps. Because oleandrin served as the reference compound in this study, we subsequently compared key pharmacological and biochemical properties of KDC203 and oleandrin side-by side (**Fig 1**).

**Fig 1.**
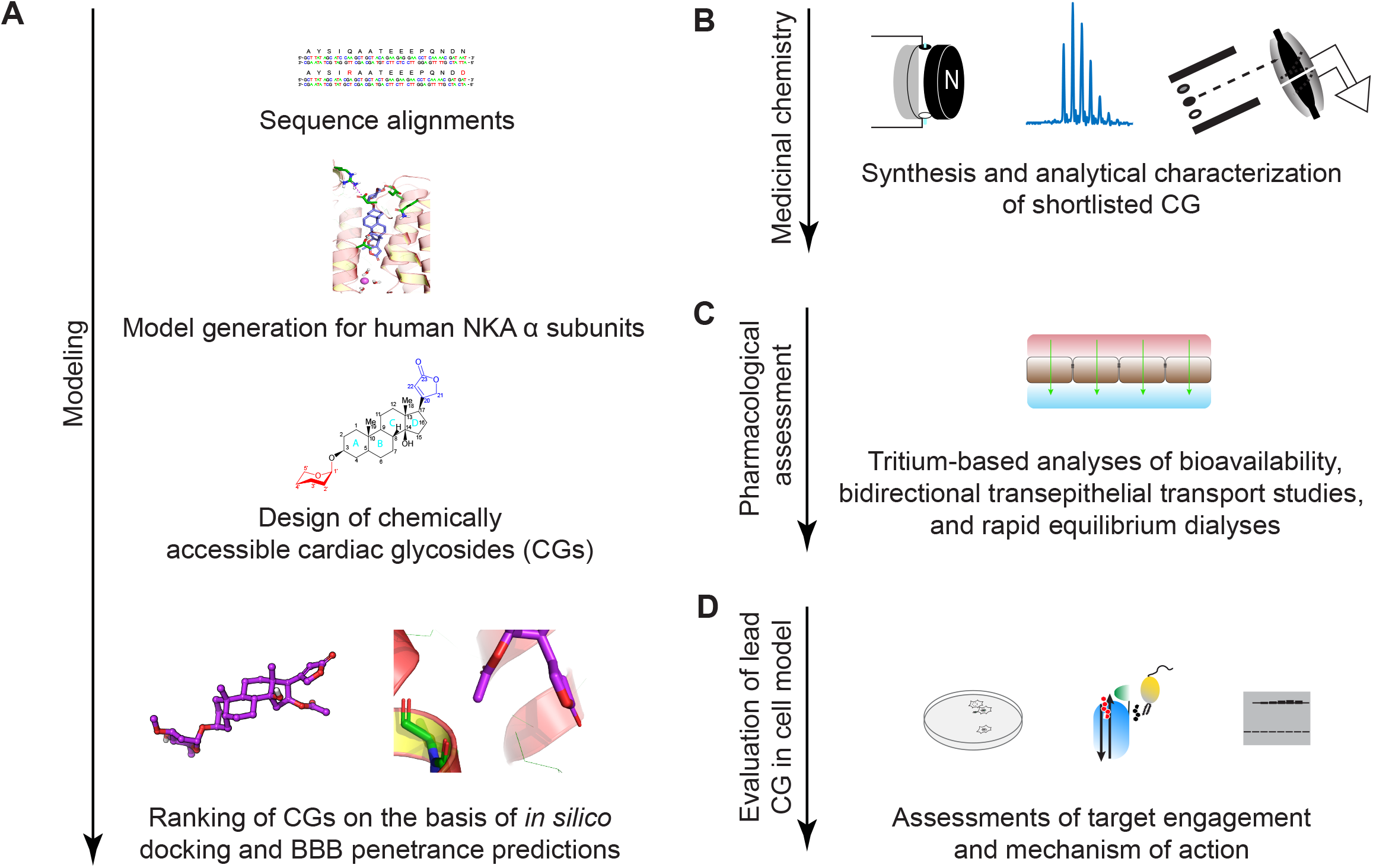
Study design. (A) *In silico* identification of a CG with favorable predicted docking characteristics and BBB barrier penetrance. (B) Synthesis and characterization of short-listed CG. (C) Assessment of BBB penetrance and distribution in relevant tissue. (D) *In vitro* characterization of potency and mechanism of action for PrP^C^ reduction of lead CG in cell-based assay.

### *In silico* modeling of oleandrin binding to human NKA α subunit

To be able to compare *in silico* the NKA binding properties of CGs for which no high-resolution binding data are available, a suitable model needed to first be generated and evaluated. Several high-resolution models have been published for shark and porcine NKAs [40–43]. These show CGs to occupy a binding pocket that can be accessed from the extracellular space and is molded from amino acid residues contributed by the α subunit (**Fig 2A**). Importantly, different CGs show highly conserved binding to this site even in NKA structures captured in different ion binding states (**Fig 2B**). To assess if building any human NKA model for the intended objective based on the porcine structures would be meaningful, we initially compared the sequence similarity between human and porcine NKA α subunits (**Fig 2C**), which indicated almost perfect sequence identity amongst ATP1A1 orthologs and showed that even human ATP1A2 and ATP1A3 paralogs share 87% sequence identity to porcine Atp1a1. Critically, the alignment of amino acid residues known to line the CG binding pocket in porcine Atp1a1 with the homologous human residues, predicted that key properties of the binding pocket are maintained in human α subunits, including the existence of previously described hydrophobic and charged internal faces (**Fig 2D**). Assured that model building based on the porcine structure is likely to provide relevant predictions for the human NKA α subunits, we focused our modeling on ATP1A1 and ATP1A3. These human NKA α subunits are expressed in brain neurons, and unless models can be built for them, the whole undertaking might be futile. To this end, we applied docking algorithms to predict binding poses for oleandrin and to evaluate the likelihood of binding poses to exist naturally based on the free binding energies (ΔG) associated with them. Two main methods were used to evaluate the proposed ligand-protein complexes, the generalized Born model (**Fig 2E**) and the slower but more accurate Molecular Mechanics-Poisson Boltzmann Surface Area (MM-PBSA) method, which computes the free binding energy by subtracting the free energy of the docked ligand-receptor complex from the free energies of its separate components [44] (**Fig 2F**). The latter revealed a free binding energy of −62.56 kcal/mol for a surface area-optimized binding pose of oleandrin within the canonical CG binding pocket of ATP1A3. The subsequent comparison of this binding pose revealed exquisite structural alignment with the experimentally observed ouabain binding pose, validating it to be a highly plausible naturally occurring pose (**Fig 2G**). A parallel screen of the pertinent literature for information on structure-activity relationship (SAR) data and features that may increase the brain bioavailability of CGs (**Fig 2H**) directed our attention toward the CG sugar moiety and opportunities to derivatize C16 within the steroid core.

**Fig 2.**
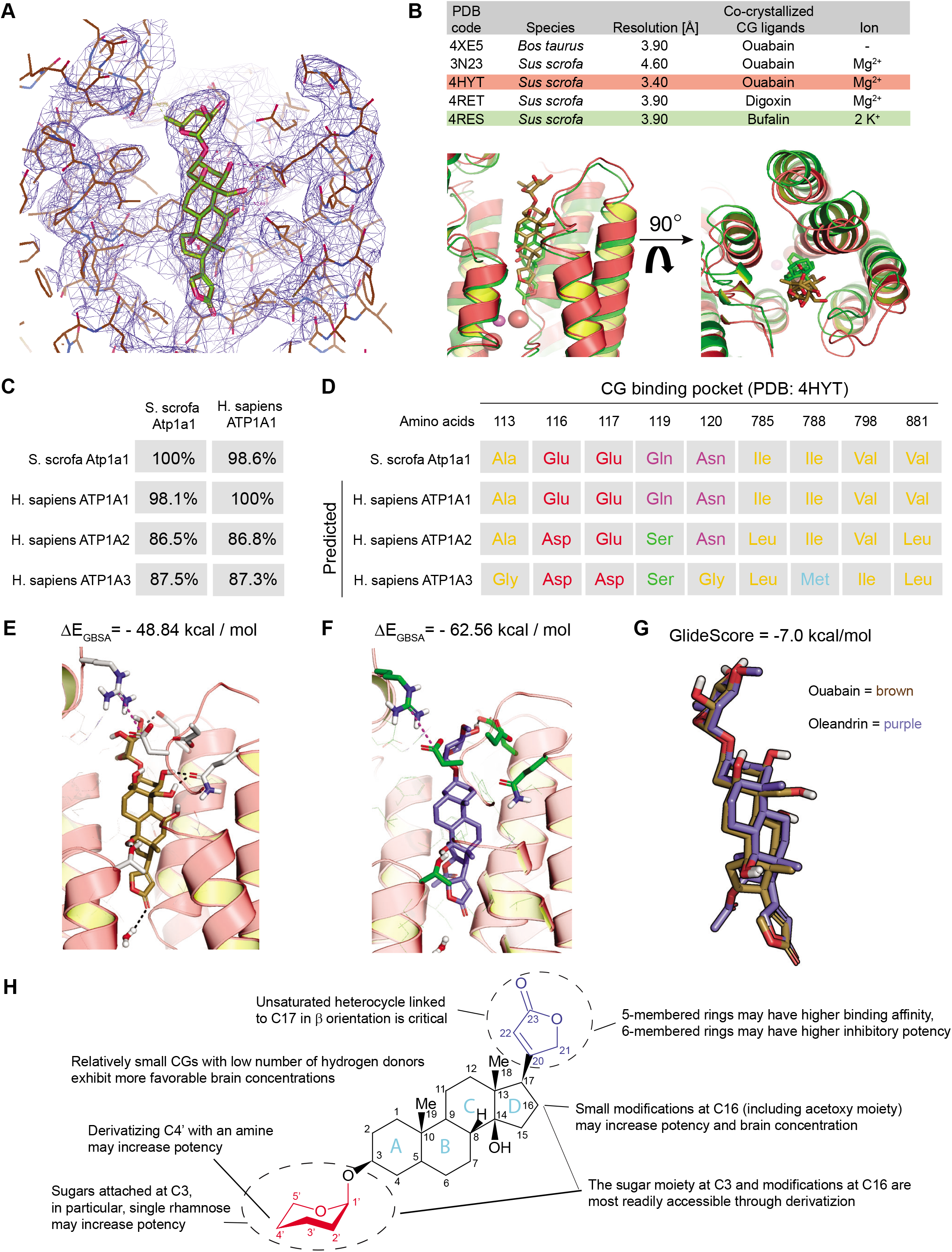
*In silico* prediction of CG binding pose within relevant human NKA α subunits and summary of key CG design considerations. (A) Electron densities of CG binding pose in PDB entry 4HYT [41]. (B) Structural alignment of ouabain (brown) and bufalin (green) within the CG binding pocket of two well-resolved X-ray crystallography models (PDB entries 4HYT and 4RES [69]) of NKAs. (C) Sequence identities between porcine and human NKA α subunits known to be expressed in the brain. (D) Comparison of observed porcine and predicted human amino acids lining the CG binding pocket. (E) Generalized Born binding pose and predicted free binding energies for oleandrin within human ATP1A3 model. (F) Surface area-optimized binding pose and predicted free binding energies for oleandrin within human ATP1A3 model. (G) Spatial alignment of observed and predicted binding poses of ouabain and oleandrin in experimentally deduced ATP1A1 and predicted surface-area optimized ATP1A3 models, respectively. (H) Key considerations for the design of potent CGs with optimized brain bioavailability drawn from structure-activity relationship data and medicinal chemistry considerations.

### Evaluation of chemically accessible oleandrin derivatives based on binding score and brain bioavailability

Our subsequent efforts focused on 270 CGs that we anticipated to be chemically accessible through combinatorial derivatization of five CG scaffolds (**Fig 3A**). Rather than synthesizing these CGs, we decided to shortlist molecules in this compound library that were predicted to have promising binding characteristics *and* improved brain bioavailability (relative to oleandrin) through a virtual screen. Initially, compounds were evaluated on the basis of physicochemical properties that determine the propensity for brain penetrance, which can be computed as a multi-parameter optimized (Version 2) score, first introduced by Zoran Rankovic [45] (**Fig 3B**). Once the protonation state and partial charges were considered, the number of virtual screen entries increased to 330. Using our human ATP1A3 model and the MM-PBSA docking method, these entries gave rise to 815 binding poses, a majority of which could be dismissed due to unfavorable internal energies or docking to a site that deviated from the canonical CG binding site. Data obtained when evaluating the 20 CG permutations defined in Scaffold 1 illustrate a typical result (**Fig 3C**). In this case, six of the 20 CGs were rejected due to excessive internal energies, or a binding pose that deviated fundamentally from the canonical CG binding pose. Rankovic MPO.v2 scores assigned to the remaining 14 CGs associated with this scaffold ranged between 2.33 and 3.00 and therefore provided no improvement of predicted brain penetrance relative to oleandrin. Finally, Glide docking scores for these CGs ranged from −4.60 to −7.35, indicating that a subset of these CGs were predicted to have improved free binding energies, relative to oleandrin.

**Fig 3.**
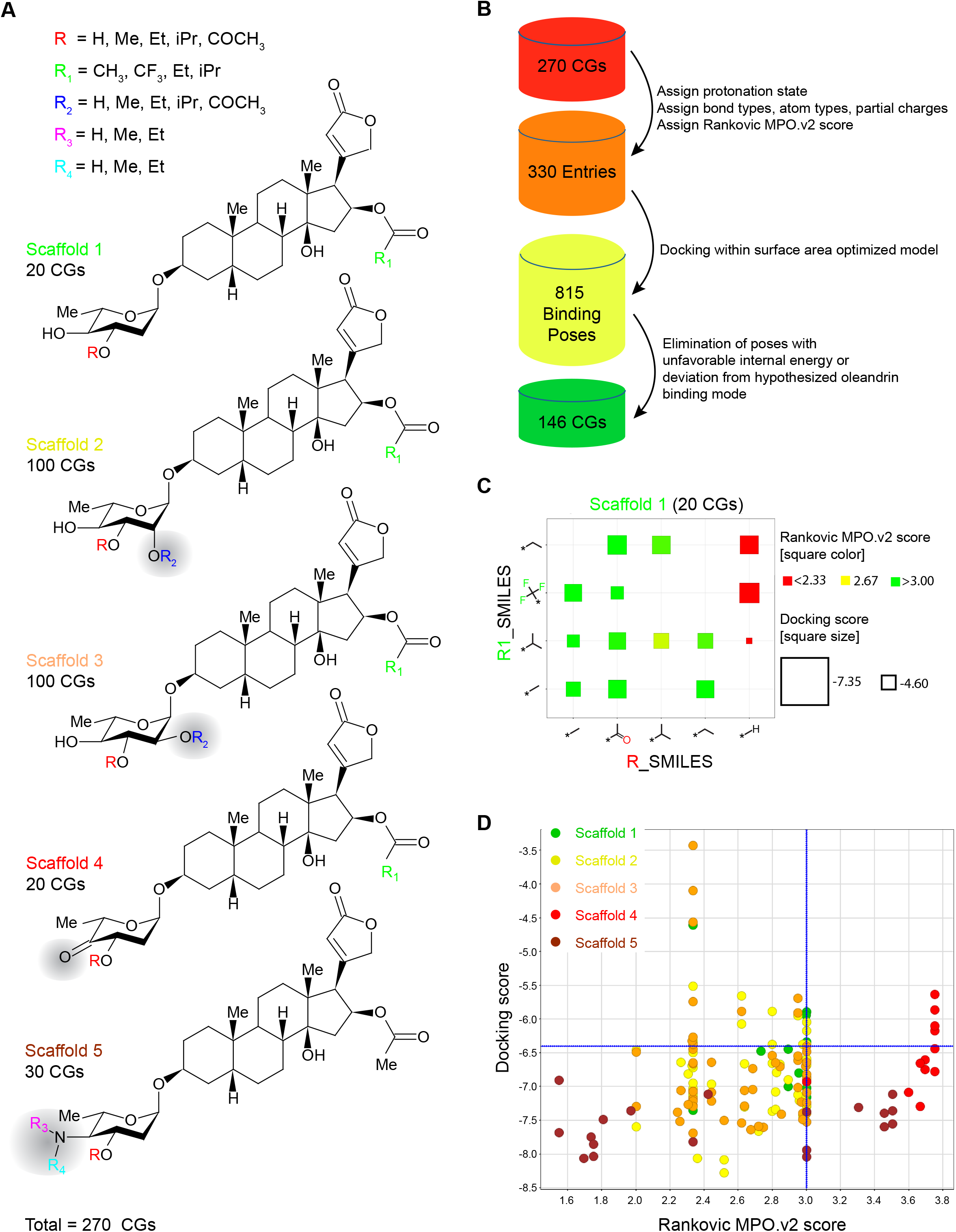
*In silico* filtering of CG derivatives accessible through chemical derivatization based on their predicted docking scores and Rankovic multi-parameter optimized (MPO, version 2) BBB penetrance scores. (A) Design of CG scaffolds and chemically accessible modifications evaluated, giving rise to a combinatorial total of 270 CGs considered. (B) Workflow for the *in-silico* assessment of candidates. The assignment of protonation states and partial charges increased total CG ligands evaluated to 330, a number that further increased to 850 once alternative binding poses, predominantly caused by freely rotating bonds were considered. The elimination of binding poses with unfavorable internal energy or deviation from the hypothetical oleandrin binding pose led to a 146 CGs that passed these filters (C) Exemplary chart depicting results from the evaluation of Scaffold 1 CG derivatives. The color scheme reflects Rankovic MPO.v2 scores [45] (a high score, represented by a green square, indicates high predicted BBB penetrance) and the size of squares reflects docking strength (a low docking score, represented by a large square, indicates strong binding). The absence of a square indicates that no binding pose that passed filter criteria (see above) was found. (D) Summary chart depicting results from the evaluation of Rankovic MPO.v2 scores and docking scores for all five scaffolds. Note that CGs derived from Scaffold 4 had similar docking scores as derivatives from other scaffolds but excelled based on their high predicted BBB penetrance.

Overall, 146 CGs passed the filters we had applied. Amongst these, Scaffold 4-derived CGs were remarkable because they exhibited the widest breadth of Rankovic MPO.v2 scores and were consistently predicted to improve binding to ATP1A3. In contrast, all Scaffold 5-derived CGs were predicted to exhibit improved brain penetrance but only a subset of these exhibited improved free binding energy relative to oleandrin (**Fig 3D**).

### Synthesis of lead compound with favorable characteristics

Results from the virtual screen recommended a small number of molecules for further analyses on the basis that they were predicted to exhibit comparable or improved binding to human ATP1A3 *and* to pass the BBB better than the oleandrin reference compound, which was computed to have a Glide docking score of −6.4 and an intermediate Rankovic MPO.v2 score of 3.0 (**Fig 4A**). More specifically, we shortlisted eight molecules, KDC201 to KDC208 (**Fig 4B, C**), prioritizing KDC203 (4’-dehydro-oleandrin) for this study because (i) it can be easily obtained through derivatization of oleandrin [46], (ii) constitutes an intermediate for the synthesis of KDC201-KDC206, and (iii) was predicted to have a high Rankovic MPO.v2 score of 3.75. Although not pursued here, replacing the C4’ hydroxyl with an amine is expected to further increase binding to ATP1A3 (KDC207) (**Fig 4D**). Similarly, introducing a C2’ methoxy group (KDC201) or a C3’ isopropyl moiety (KDC205) within the KDC203 sugar were predicted to further improve binding to ATP1A3 (**Fig 4E**). Finally, a gain in the Glide docking score was consistently attained when we replaced hydrogen atoms within the C16 acetoxy group of the oleandrigenin core with fluorine atoms (KDC202, KDC204, KDC206, KDC208) (**Fig 4F**). To move from *in silico* to biochemical analyses, we obtained KDC203 through oxidization of oleandrin in dichloromethane followed by HPLC clean-up as a white solid with >95% purity (assessed by ^1^H NMR). A stably tritiated version of this compound was obtained commercially through a customized labeling request.

**Fig 4.**
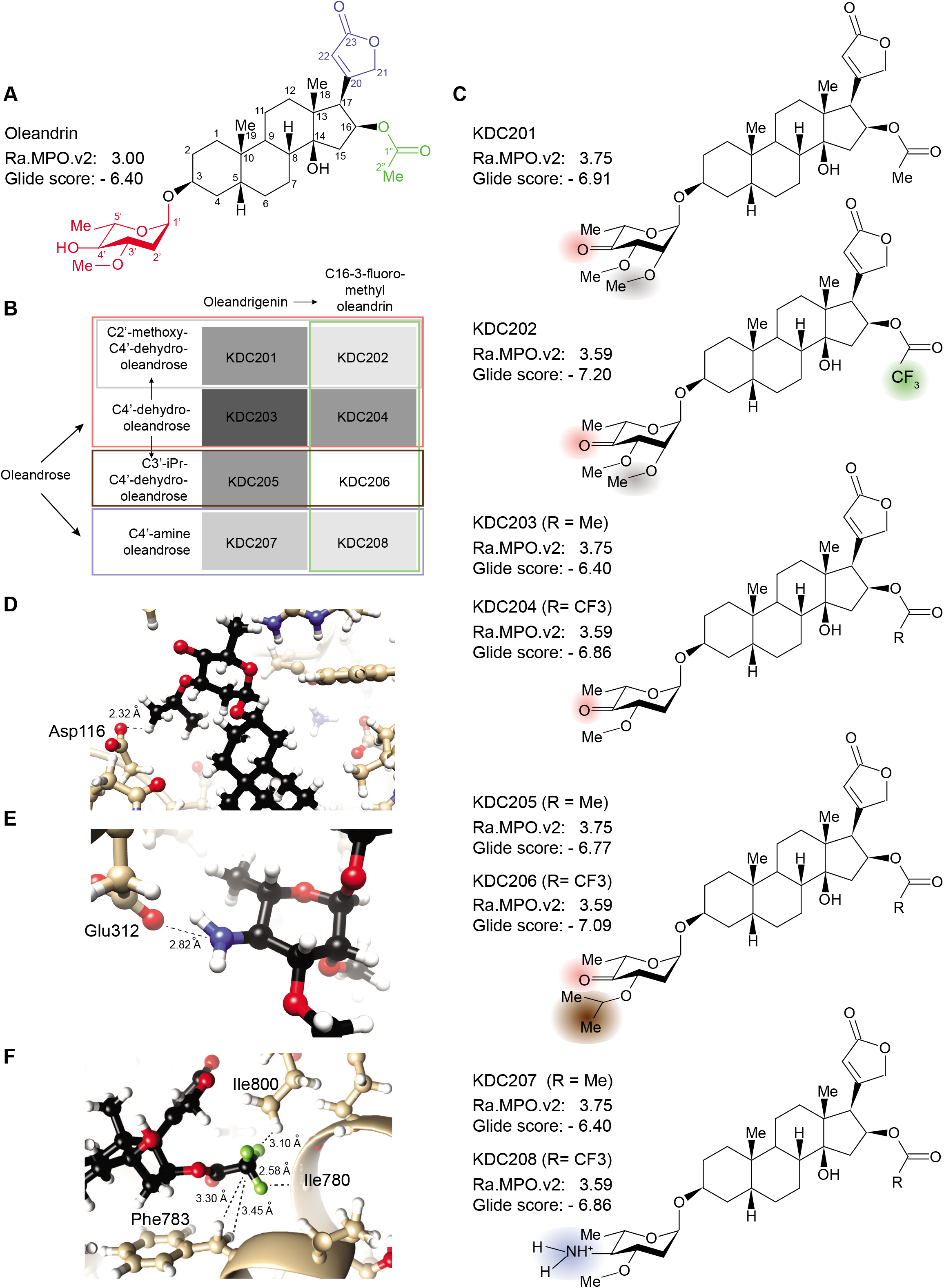
Features of chemically accessible Scaffold 4 derivatives-of-interest. (A) Chemical structure and reference Rankovic MPO.v2 and docking scores of oleandrin. (B) oleandrin derivatives of interest that should be chemically accessible with good yields in few steps. Starting with oleandrin, several compounds with high predicted Rankovic MPO.v2 scores can be accessed through C4’-dehydro-oleandrin (here termed KDC203) or C4’-amine-oleandrin [46]. (C) Chemical structures of compounds listed in Panel B with their respective Rankovic MPO.v2 and Glide scores. Note that structures of KDC204, KDC206 and KDC208 are not depicted. (D-F) Close-up models providing plausible explanations for predicted increases of Glide docking scores (residue numbers as in PDB 4HYT). (D) Salt bridge between the C4’ amine and the sidechain of glutamic acid residue 312. (E) Hydrogen bonding opportunity for the C3’ isopropyl group with the side chain of aspartic acid residue 116. (F) Good fit of C16 trifluoro-acetoxy group within hydrophobic binding pocket shaped from side-chain atoms of phenylalanine 783 and isoleucine 800 residues lining the CG binding pocket.

### Analysis of brain bioavailability

To assess the bioavailability of KDC203 in a model whose NKAs respond similarly to CGs as their human orthologs, wild-type rodents would not be a good choice, because they have long been known to exhibit more than hundred-fold lower sensitivity toward a large range of CGs than most other mammals [47–49], a difference that has since been attributed to specific differences in Atp1a1 amino acids lining the CG binding pocket. Rather than moving to larger mammals, whose ATP1A1-genes may be more similar to the human ortholog in this regard, we pursued these studies with Atp1a1 gene-edited mice engineered to carry two point mutations in the first extracellular loop, namely amino acid exchanges Q111R and N122D (amino acid numbering as for the PDB entry 4HYT) coded by the Atp1a1 gene (Atp1a1^S/S^) [50]. These point mutations are understood to sensitize this α subunit toward CG binding by making it more “human-like” [51, 52]. One day following the subcutaneous injection of five Atp1a1^S/S^ mice per cohort with tritiated oleandrin or KDC203, the mice were sacrificed, transcardiac perfused with phosphate-buffered saline (PBS) and radioisotope levels determined in their brains, hearts, kidneys, and livers (**Fig 5A**). These analyses revealed that KDC203 achieved significantly higher brain levels than oleandrin. Notably, a mean total concentration of 30.46 nM tritiated KDC203 levels was observed in the brains of five mice, a value that exceeded KDC203 levels in the heart 3.5-fold, and even surpassed mean kidney and liver levels. Taken together, this experiment established KDC203 to exhibit, relative to oleandrin, the improved brain bioavailability that its higher Rankovic MPO.v2 score (3.75 for KDC203 versus 3.0 for oleandrin) had predicted.

**Fig 5.**
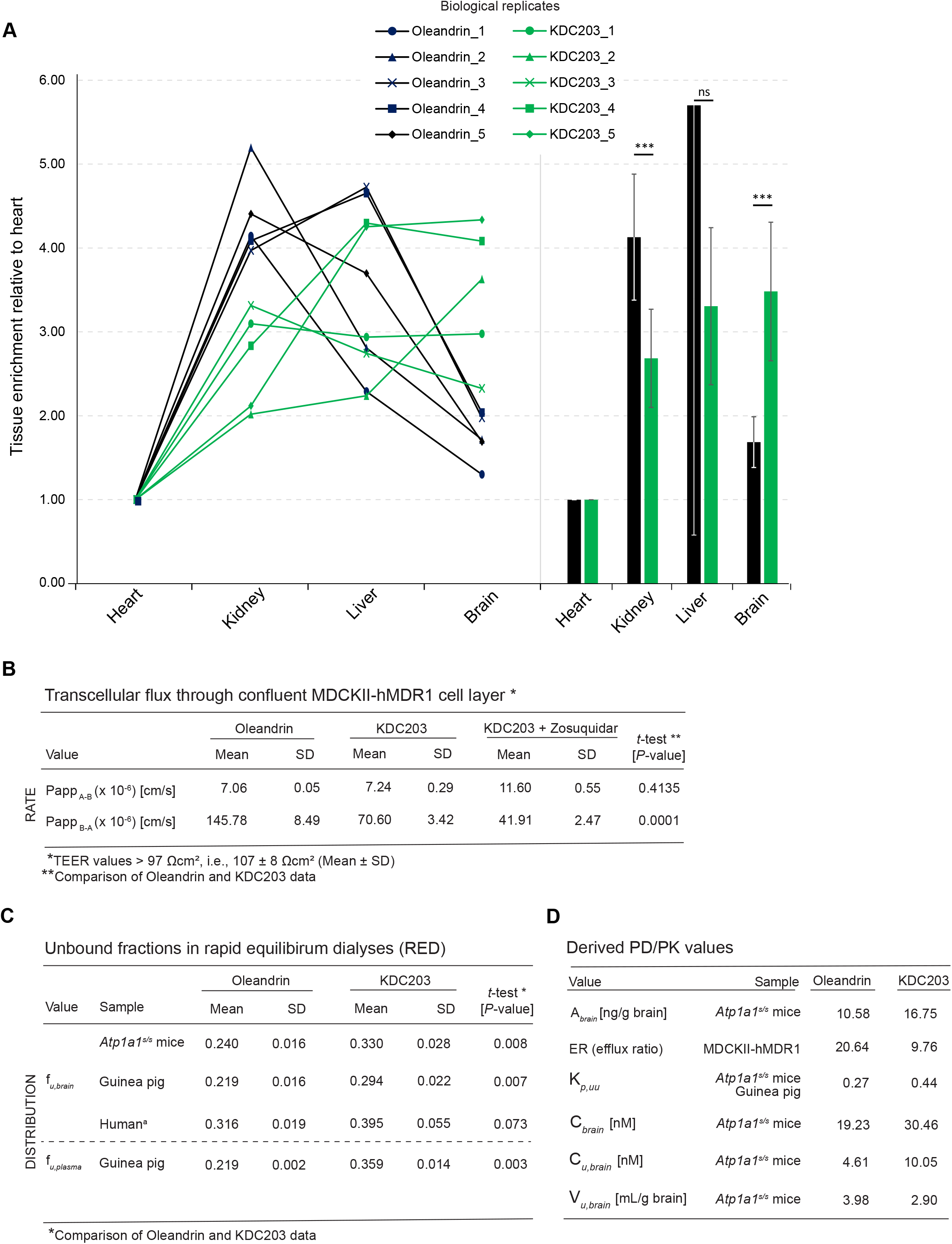
Improved brain bioavailability of KDC203 relative to oleandrin. (A) Oleandrin enriches in kidney and liver tissue and its brain and heart levels are similar 24 hours following its acute subcutaneous injection as a tritium-labeled compounds into cohorts of five mice. In contrast, KDC203 reaches highest levels in the brain and its brain levels exceeded its heart levels 3.5-fold 24 hours after subcutaneous injection. (B) KDC203 is a lesser hMDR1 substrate than oleandrin. Recorded values of apparent permeability (Papp) from apical to basolateral (A-B) and vice versa (B-A). The efflux ratio is calculated as Papp_B-A_ / Papp_A-B_. Zosuquidar was deployed as a selective inhibitor of MDR1. Measurements of the TEER on the day of experiment (day 5) resulted in a mean value of 107 ± 8 Ωcm² (± SD, n=22). Lucifer Yellow Permeability data (n=22) were generated prior to the bidirectional assay with oleandrin and KDC203 (n=3). (C) Rapid equilibrium dialyses of brain and plasma samples establish that KDC203 has a lower propensity than oleandrin to associate nonspecifically with components in the respective extracts, predictive of a higher concentration of free KDC203 that is available for specific engagement with its NKA target. (D) Pertinent pharmacological characteristics of oleandrin and KDC203 computed based on their *in vivo* tissue and plasma concentrations and the *in vitro* measurement of their unbound fractions in rapid equilibrium dialyses.

### Transepithelial diffusion and transport

To compare the rate of transcellular passive diffusion and active transport of KDC203 and oleandrin, we made use of canine MDCK epithelial cells, which are known to provide a high level of monolayer integrity through tight junctions limiting paracellular diffusion. More specifically, a second generation MDCKII cell line was employed in these experiments that had been generated by deleting the canine MDR-1 transporter and replacing it with the human MDR1 gene. The permeability screen was conducted as a bidirectional assay. Thus, first, the rate of transepithelial flow from apical to basolateral (Papp_A-B_) was recorded, revealing a slightly lower value of 7.06 x 10^-6^ for oleandrin than KDC203, which crossed the epithelium at a rate of 7.24 x 10^-6^ cm/s (**Fig 5B),** with both compounds showing a considerably higher permeability than published values of 0.5 x 10^-6^ cm/s for digoxin [53, 54]. A striking difference between oleandrin and KDC203 became apparent when measuring the reverse rate of transport from basolateral to apical (Papp_B-A_), which indicated that oleandrin was being actively extruded at 145.79 x 10^-6^ cm/s, more than twice the 70.60 x 10^-6^ cm/s rate of KDC203 extrusion. Consequently, efflux ratios (ERs), which are computed as the ratio of the rates of extrusion versus intrusion, exceeded for both CGs values of 2, thereby establishing both CGs as hMDR1 substrates. Consistently, when we next added the MDR-1 inhibitor Zosuquidar, the extrusion of KDC203 was reduced by more than 50%, further corroborating its hMDR1-dependent extrusion.

### Free versus bound fraction

Next, we measured the free unbound (f_u_) concentrations of oleandrin and KDC203 in rapid equilibrium dialysis (RED) experiments. Deploying brain homogenates of *Atp1a1^S/S^* mice, guinea pigs, or humans, we observed that the f_u,brain_ values of oleandrin were lower than the corresponding values for KDC203, indicative of lower unspecific binding of KDC203 to biological surfaces within the brain (**Fig 5C**). This difference reached robust significance (*P* < 0.005) when studying brain homogenates of *Atp1a1^S/S^* mice and guinea pigs and barely missed significance with human brain homogenates.

From these experimental results, additional pharmacokinetic characteristics of oleandrin and KDC203 could be deduced (**Fig 5D**). KDC203 exhibiting a twofold lower efflux ratio than oleandrin predicted an improvement in its CNS exposure that is consistent with the experimentally determined increase in its total brain levels (**Fig 5A**). Calculations of K_p,uu_ values for both oleandrin (0.27) and KDC203 (0.44) resulted in values below 1, thereby revealing possible efflux processes at the BBB, with oleandrin again exhibiting a higher degree of transport asymmetry. V_u,brain_ values for oleandrin and KDC203 of 3.98 mL/g and 2.90 mL/g brain, respectively, express the tendencies of the compounds to reside outside of the interstitial fluid (ISF), e.g., bound to brain tissue. Finally, the free mean concentration of these CGs in the brain, expressed as *C_u,brain_*, that is influenced by the *extent* of CNS-delivery as well as its intracerebral *distribution* computed to 4.61 nM for oleandrin and 10.05 nM for KDC203, thereby predicting the latter to reach a more than twofold higher unbound concentration level in the brain.

### *In vitro* assessment of potency in differentiated ReN VM cells

Because the *in silico* docking analyses of oleandrin and KDC203 had returned identical Glide scores of −6.40 (**Fig 4**), we anticipated that both compounds may exhibit similar potencies in an assay that relies on the engagement of these compounds with their cognate NKA α subunit binding site. To evaluate this characteristic experimentally in a relevant model, immortalized human cells (ReN VM cells) that we had differentiated to acquire neural or astrocytic characteristics were exposed for a duration of one week to low nanomolar levels of oleandrin or KDC203. Following cell lysis, steady-state ATP1A1 levels were assessed by western blot analysis to determine if the ligand-receptor interactions had caused the previously observed reduction in ATP1A1 levels [18]. This analysis established that KDC203 had indeed similar potency as oleandrin on the basis that both drugs achieved a similar reduction in ATP1A1 levels when added at 4 nM concentration to the cell culture medium (**Fig 6A**). However, whereas oleandrin was toxic at concentrations exceeding 4 nM [18], KDC203 was tolerated by the cells at all concentrations tested in this initial pilot experiment. Remarkably, exposure of ReN VM cells for one week to 8 or 16 nM levels of this compound led to additional reductions in ATP1A1 levels. Further analyses in ReN VM cells established that the reduction in the levels of the NKA α subunit ATP1A1 is paralleled by a lesser yet also pronounced reduction in the levels of the NKA β subunit ATP1B1 (**Fig 6B**). As for steady-state PrP^C^ levels, the quantitation of intensities of western blot bands revealed that a maximum 84% reduction in 3F4-reactive signals was attained when ReN VM cells were exposed for one week to 12 nM concentrations of KDC203 (**Fig 6C**). To assess whether the KDC203-dependent reductions in the steady-state protein levels of ATP1A1 and PrP^C^ represent an idiosyncrasy of ReN VM cells or can also be observed in another human neural cell model, we next repeated the analysis with differentiated T98G cells. Noticing that the KDC203-dependent reduction in ATP1A1 was less pronounced in T98G than in ReN VM cells, we extended the western blot analyses to other NKA α subunits that are known to be expressed in human brains. Remarkably, not only did this cell model recapitulate the KDC203-dependent reduction in PrP^C^ levels but it attained it without exhibiting a similarly profound diminution of the three NKA α subunits ATP1A1, ATP1A2, ATP1A3 (**Fig 6D**).

**Fig 6.**
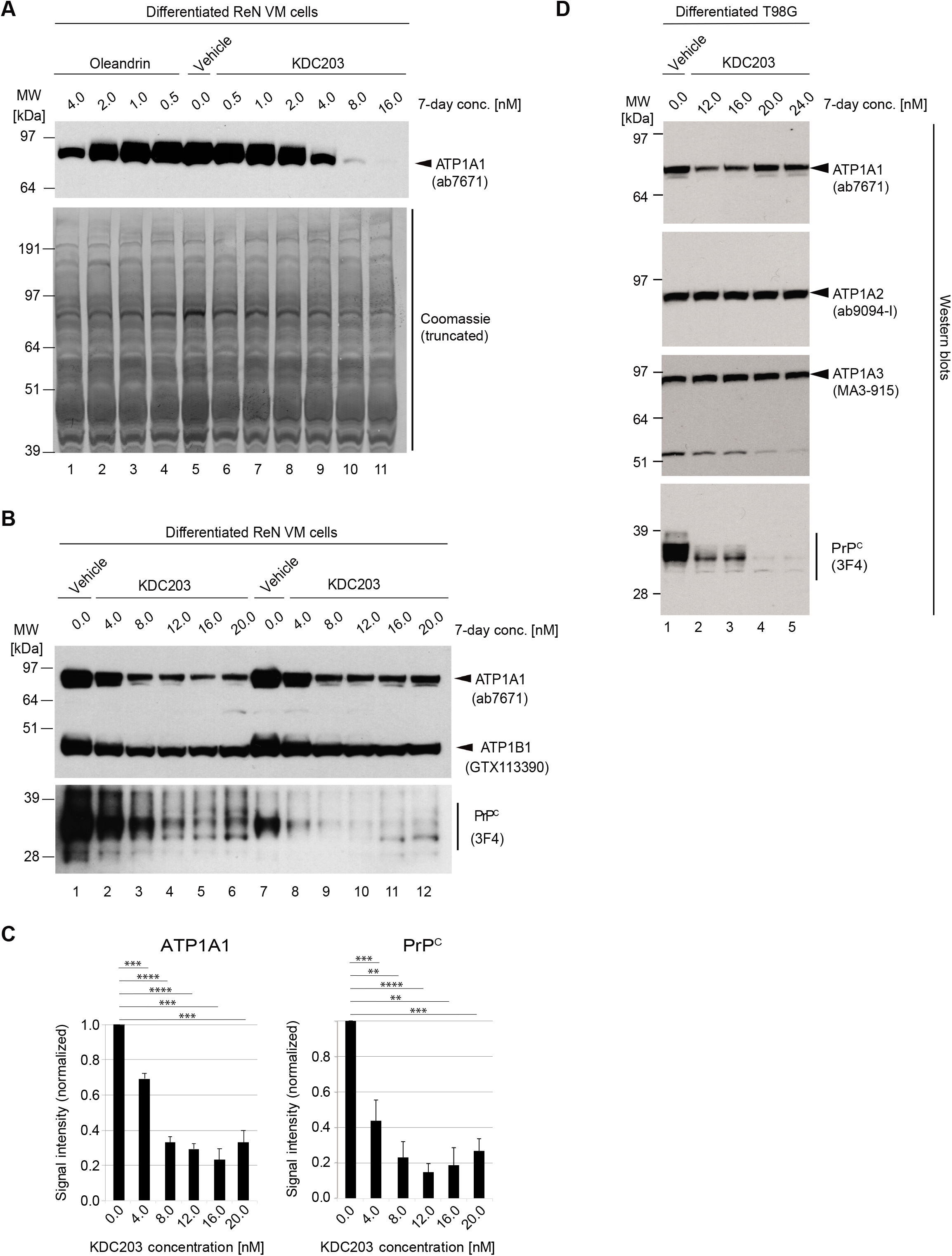
KDC203 reduces steady-state PrP^C^ levels in human neural cell lines. (A) Oleandrin and KDC203 reduce steady-state ATP1A1 protein levels to a similar degree. Side-by-side western blot-based comparison of ATP1A1 signal intensities in cellular extracts derived from ReN VM cells following 7-day treatment with the respective CGs. (B) 7-day exposure of ReN VM cells does not only affect ATP1A1 protein levels by also causes a concentration-dependent reduction in the steady-state protein levels of the NKA β subunit ATP1B1 and PrP^C^. (C) Quantitation of western blot signal intensities of ATP1A1 and PrP^C^ following 7-day KDC203 treatment of ReN VM cells at concentrations indicated, with each value being computed from the analysis of three biological replicates. (D) The KDC203-dependent reduction in steady-state PrP^C^ protein levels is not an idiosyncrasy of ReN VM cells but can also be observed in other neural cell models, including differentiated human glioblastoma cells (T98G).

### KDC203 is less toxic than oleandrin

Considering the critical role NKAs play in maintaining the electrochemical gradient in all eukaryotic cells, we wondered whether the KDC203-dependent diminution in steady-state ATP1A1 levels that we had observed in ReN VM cells was compensated for by an upregulation of ATP1A2 and/or ATP1A3. To address this point, a repeat KDC203 treatment of differentiated ReN VM cells was undertaken, this time exposing cells to up to 40 nM concentrations of the compound, and cellular extracts were again analyzed for all three NKA α subunits expected to be expressed in these cells (**Fig 7A**). Western blot results from this experiment corroborated the robust KDC203-dependent reduction in steady-state levels of ATP1A1 but also revealed that this effect of the compound extends to ATP1A2 levels. More specifically, ATP1A1 levels were diminished in cells treated with KDC203 up to 16 nM levels yet were observed to rebound slightly when KDC203 levels were further increased. In contrast, steady-state ATP1A2 levels declined as levels of KDC203 were increased, with no sign of a rebound at any concentration assessed. Intriguingly, ATP1A3 levels stayed unchanged in the presence of up to 12 nM KDC203 levels, then increased and reached a maximum in the presence of 20-32 nM KDC203 before declining again at 40 nM KDC203 levels.

**Fig 7.**
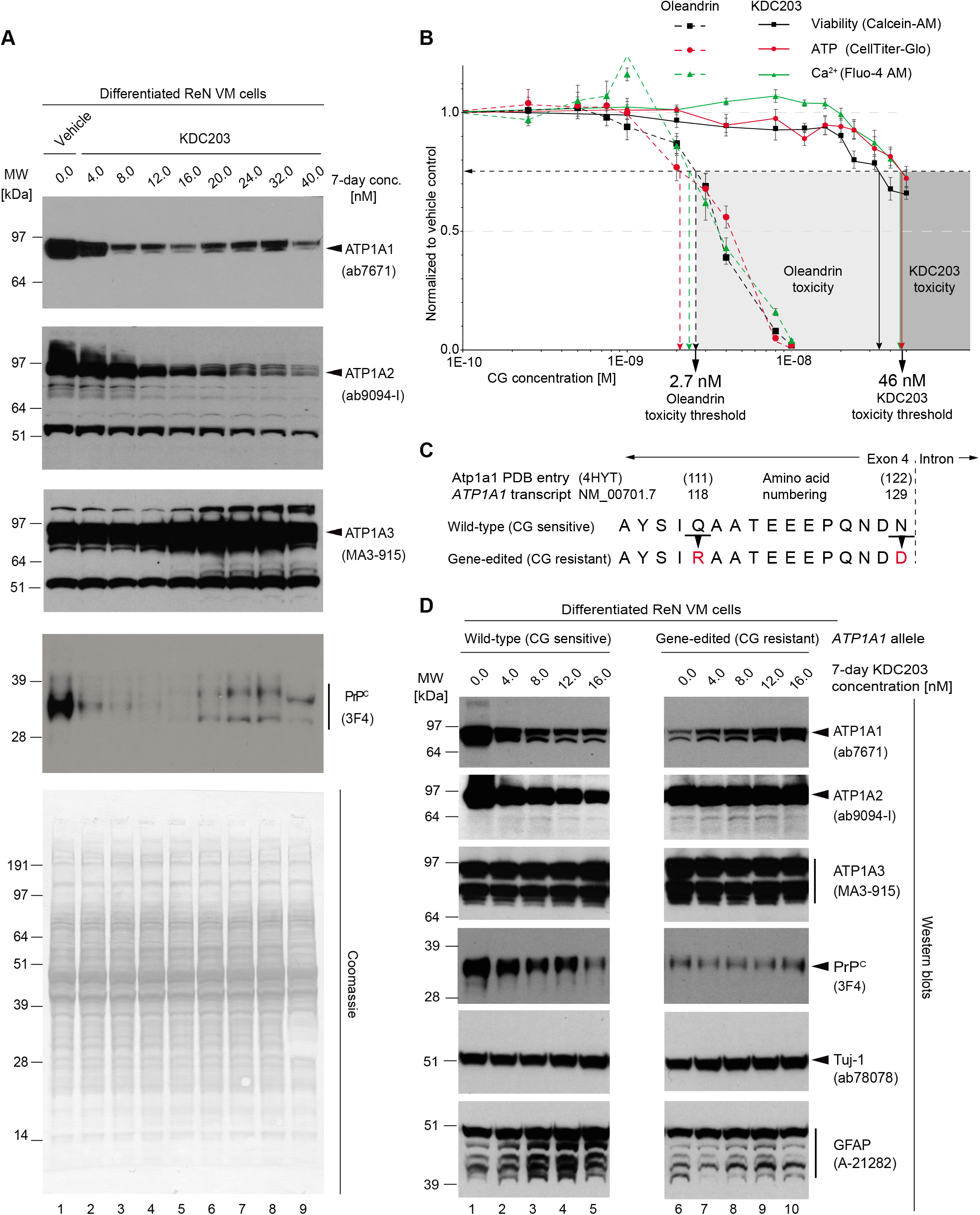
KDC203 induces a compensatory upregulation of ATP1A3, is less toxic than oleandrin, and requires binding to CG binding pocket within ATP1A1 for reducing PrP^C^ levels. (A) The KDC203 concentration-dependent reduction in the steady-state protein levels of ATP1A1, ATP1A2, and PrP^C^ is paralleled by an increase in ATP1A3 levels. Total concentrations of cellular extracts were adjusted, and equal volumes of these adjusted extracts were loaded onto the gel as evidenced by the Coomassie stain. Please note the slight rebound in ATP1A1 signals and the splitting of the 3F4-reactive signals into two bands that migrated slower and faster than the corresponding band in vehicle-treated differentiated ReN VM cells exposed to 20-32 nM KDC2039 concentrations. (B) KDC203 exhibits tenfold lower toxicity than oleandrin in three assays that were used to monitor the metabolic health of 7-day differentiated ReN VM cells. Each data point represents the mean of six biological replicates. (C) Amino acid sequence alignment of an NKA α subunit segment contributing to CG binding in wild-type human ATP1A1 protein and its mutated derivative rendered refractory to CG binding by replacing human residues 118 and 129 (numbering based on human *ATP1A1* transcript NM_00701.7) with the corresponding mouse Atp1a1 residues. (D) KDC203 causes the expected reduction in the steady-state levels of ATP1A1 and ATP1A2 in differentiated wild-type ReN VM cells but not in the gene-edited ReN VM *ATP1A^R/R^* cells whose ATP1A1 protein does not bind CGs. Despite the decrease in ATP1A2 levels, the NKA α subunit that is predominantly expressed in astrocytes, no reduction in the steady-state levels of astrocytic GFAP is observed.

When western blots from the same cellular extracts were probed with an antibody that detects human PrP^C^, its steady-state levels were again revealed to be dramatically reduced in cells exposed to 4 - 16 nM KDC203. Interestingly, in cells exposed to KDC203 concentrations of 20-32 nM, a slight rebound in the intensity of 3F4-reactive signals was visible. However, the PrP^C^ signals observed under these circumstances were split into two signals that migrated with apparent molecular weights of 30 and 35 kDa, in contrast to the dominant 3F4-reactive signal detected at 32 kDa in naïve ReN VM cells. Importantly, the KDC203 exposure of the cells did not affect bulk protein levels as these were constant for all concentrations tested. These experiments suggest that the ReN VM cell cultures were able to cope with KDC203 concentrations exceeding 8 nM by increasing their steady-state ATP1A3 levels.

Next, we interrogated the health of ReN VM cells exposed to oleandrin versus KDC203 by monitoring their metabolic activity, ATP levels, and intracellular calcium levels using fluorescent indicators (**Fig 7B**). Setting an arbitrary toxicity threshold that marked the concentration at which all three cellular health indicators dropped to 75% of levels observed in untreated cells, these analyses determined oleandrin to reach this threshold at 2.7 nM versus KDC203 at 46 nM concentrations. Taken together, these experiments corroborated the conclusion that ReN VM cells are better equipped to tolerate double-digit nanomolar concentrations of KDC203 than oleandrin and that a compensatory upregulation of ATP1A3 may play a role in the tolerance toward KDC203.

### Mutagenesis of CG binding site within ATP1A1 abolishes KDC203-dependent PrP^C^ reduction

Intrigued by our observation that KDC203 suppressed steady-state protein levels of ATP1A1 and ATP1A2 yet increased ATP1A3 levels when ReN VM cells were exposed to concentrations exceeding 16 nM, we wondered if the levels of these α subunits were independently or interdependently affected by KDC203. We had previously observed that the CG-dependent reduction in the steady-state protein levels of ATP1A1 and PrP^C^ could be rescued by rendering the human ATP1A1 protein resistant to high-affinity CG docking [18], thereby establishing the need for CG and NKAs forming a ligand-receptor interaction to achieve this outcome, as opposed to other, less defined effects of CG on the cell. To obtain this result we had mutated the *ATP1A1* allele in two positions such that two amino acid residues known to contribute to CG binding are instead coding for the corresponding amino acids present in mouse Atp1a1 [55] (**Fig 7C**). Here, we made use of this ReN VM-derived *ATP1A1^r/r^* cell model to investigate whether the KDC203-induced changes to ATP1A2 and ATP1A3 are triggered by KDC203 binding directly to the respective cognate binding sites in these NKA α subunit paralogs or are also caused by KDC203 docking to ATP1A1 and affecting ATP1A2 and ATP1A3 indirectly. Remarkably, abrogating binding of KDC203 to ATP1A1^r/r^ not only precluded the changes in steady-state levels of ATP1A1 but also prevented changes to the steady-state levels of ATP1A2 and ATP1A3 (**Fig 7D**). Moreover, these experiments validated the hypothesis that the KDC203-mediated reduction in steady-state levels of PrP^C^ depends on this CG predominantly forming a ligand receptor complex with ATP1A1 because no KDC203-mediated reduction in PrP^C^ levels was observed in the ATP1A1^r/r^ cell model. Considering the well-known preferential expression of ATP1A2 and ATP1A3 in astrocytes and neurons, respectively, we next wondered if the KDC203-depedent reduction in ATP1A2 and concomitant increase in ATP1A3 steady-state levels reflected a shift to a more neuronal differentiation state. To this end, we compared levels of the neuron-specific class III β tubulin (Tuj-1) and the astrocytic glial fibrillary acidic protein (GFAP) in ReN VM cells treated with KDC203. Western blot analyses of cellular extracts, which had been adjusted for total protein, revealed that the KDC203-mediated, ATP1A1-dependent decrease in PrP^C^ levels was not paralleled by significant changes in the levels of Tuj-1 or GFAP (**Fig 7D**). Taken together this experiment highlighted the importance of KDC203 being able to dock to its cognate binding site on ATP1A1 for all changes in steady-state levels described here. It also suggested that the KDC203-induced shift in the expression profile of NKA α subunits is indicative of a plasticity in the expression of these paralogous subunits, perhaps as part of a compensatory rescue for the loss of ATP1A1, rather than representing a facet of cells undergoing broad astrocyte-to-neuron reprogramming in the presence of KDC203.

## DISCUSSION

This report described a systematic approach to the identification and validation of a CG that offers improved brain bioavailability relative to other molecules within this compound class. The work presented is part of a larger research program aimed at the development of a treatment for prion diseases. It focused on pharmacological properties, potency, and mechanism of action of a lead CG, termed KDC203, that reduces PrP^C^ levels by targeting NKAs in its immediate proximity. Starting with oleandrin, a CG that has shown some promise for brain-related applications [28, 29, 56, 57], we sought to avoid resource-intensive *in vitro* screens by pairing *in silico* modeling and BBB penetrance predictions with insights into chemically accessible derivatives. Using this pragmatic approach, a chemical space of more 270 CGs was filtered to less than ten compounds whose Rankovic MPO.v2 scores and Glide scores for binding to NKAs were predicted to equate or surpass the corresponding scores for oleandrin. Subsequent pharmacological and biochemical analyses focused on KDC203, establishing that its brain bioavailability is approximately twofold improved relative to oleandrin. The compound is stable at 36.6°C for extended periods of time (**S1 Fig**), and the treatment of ReN VM cells with 12 nM concentrations of KDC203 suppressed steady-state PrP^C^ levels by as much as 84% but had no impact on the overall expression of most proteins in these cells. Our biochemical analyses indicate that at least in ReN VM cells the PrP^C^ level reduction achieved with KDC203 depends on the compound forming a ligand-receptor complex with ATP1A1, which in turn leads to a reduction in ATP1A1 and ATP1A2 levels and an increase in ATP1A3. The CRISPR-Cas9-driven replacement of two amino acids, predicted to contribute to the established CG binding site within ATP1A1, prevented all these changes to the steady-state protein levels of NKAs and PrP^C^.

The ability of certain plant and animal species to synthesize CGs represents an adaptation that serves as a defense against herbivores and predators. Because this system is ancient and predated the existence of organisms with complex brains, an efficiency to pass the BBB is unlikely to have played a role in the evolution of CGs. Instead, compounds within this class act on cells by potently breaking their electrochemical gradient, leading at least in mammals to fatal cardiac arrhythmias as the cause of death. This reality represents both a challenge and opportunity for the objective to identify CGs with improved brain bioavailability; it suggests that the failure of available CGs to pass efficiently into the brain may not reflect a fundamental challenge that even extended fitness selection was not able to solve but rather constitutes an untapped opportunity. KDC203, chemically known as 4’-dehydro-oleandrin, our current lead compound for brain CG applications is not novel. A method for its derivatization from oleandrin was first reported in a patent application from 1975 [58] as one of several CGs that emerged around that time from a larger drug development program of the German Beiersdorf AG. The original inventor described 4’-dehydro-oleandrin as a “good cardiotonic, and particularly suitable for use as a medicament in the treatment of cardiac insufficiency”. Its oral toxicity assessed in cats was almost twofold lower than the corresponding values for oleandrin, yet its oral effectiveness—a measure determined by comparing the lethal doses of a given compound following its oral versus intravenous administration—was approximately 20% higher. To our knowledge, the only other mention of this compound in the primary literature can be found in a report from 2016, which described its synthesis next to the synthesis of other C4’-substituted oleandrin derivatives and determined that its cytotoxicity was slightly reduced relative to oleandrin [46]. The authors reported an IC_50_ of 46 nM (after 72 hours exposure of cells) toward HeLa cells. In this study, we observed cytotoxicity after 7-day treatment of ReN VM cells at KDC203 concentrations upward of 40 nM.

A future prion therapeutic need to provide protection over extended periods. Consequently, the *rate* of uptake is of lesser concern than the brain concentration of a treatment compound available for target engagement and its safety profile over time. Hence, pharmacologically, the focus should be on the *extent* to which an orally administered CG accumulates in unbound form, a prerequisite for target engagement, in the brain versus other organs over time. Both the *in vitro* MDCKII assay data and the calculated K_p,uu_ values suggest that KDC203 will be subjected to active extrusion at the BBB. Encouragingly, the data presented here suggest that KDC203 is a lesser hMDR1 substrate than oleandrin and reaches within 24 hours higher levels in the brain than in other tissues we investigated, including the heart. These improved pharmacological properties of KDC203 can be rationalized by the removal of the hydroxyl group, which made this molecule more hydrophobic. A comparison of the BBB penetrance of large numbers of compounds indicated that the capacity of a given compound to form hydrogen bonds correlates inversely with their brain penetrance. In fact, aside from a compound’s molecular size no other characteristic has been shown to be more predictive of its brain penetrance [45].

The *V_u,brain_* values of oleandrin (3.98 mL/g brain) and KDC203 (2.90 mL/g brain) that we computed are in a comparable range to other CNS-therapeutics like the opioid morphine (2.1 mL/g brain) [59] or the central-acting antihypertensive drug clonidine (5.9 mL/g brain) [60]. Naturally, data from equilibrium dialyses may not accurately predict *in vivo* distributions, since the homogenization destroys biomolecular structures and, consequently, cannot capture biology that relies on intact cells, including the ‘cellular trapping’ that can at times be observed in intracellular acidified compartments for basic compounds [60].

The subcutaneous injection of identical doses of oleandrin or KDC203 (0.6 mg/kg) yielded pharmacologically active unbound brain concentrations of 4.61 nM for oleandrin and 10.05 nM for KDC203 (**Fig 5**), near the concentration of this compound (12 nM) needed to generate the most profound reduction in PrP^C^ levels *in vitro* (**Fig 6**). The distinct pharmacological properties of oleandrin and KDC203 may reflect differences in non-specific interactions with brain tissue *and* differences in their total brain levels, as indicated by their unbound drug partitioning coefficients (*K_p,uu_*). A crude estimation based on extrapolations suggests that a 10 nM concentration of unbound KDC203 might be attainable in human brains (**S2 Fig**).

An interesting facet of our results is the observation that ReN VM cells exhibit considerable plasticity and interdependency regarding the steady-state levels of their NKA α subunits. As levels of ATP1A1 decreased upon KDC203 engagement, so did ATP1A2 levels in a manner that depended on the reduction in ATP1A1 levels. In contrast, ATP1A3 levels increased together with KDC203 levels in the cell culture medium, suggesting that the levels of this α subunit are not tied to ATP1A1, and instead may offer some level of functional compensation for the loss of ATP1A1 and ATP1A2. Moreover, our data suggest that in cells the KDC203-dependent reduction in PrP^C^ levels can exceed the reduction in the levels of any of the NKA α subunits. If the most parsimonious explanation still applies here, namely that the same fundamental mechanism of action of a CG-dependent co-internalization of NKAs and PrP^C^ is responsible across cellular paradigms, this finding may reflect differences in the rate at which distinct cell lines can replenish internalized NKAs versus PrP^C^ at the cell surface.

## CONCLUSION

This work established KDC203, chemically known as 4’-dehydro-oleandrin, as a lead CG for brain applications. KDC203 exhibited, relative to oleandrin, several improvements to its pharmacological properties: it reached higher CNS concentrations, presumably because it is a lesser MDR1 substrate and exhibits lower unspecific sequestration in plasma and brain tissues. It also was less toxic than oleandrin yet exhibited similar potency in *in vitr*o PrP^C^ reduction assays. Our *in-silico* work predicted additional molecules to have favorable brain penetrance. A subset of these can be easily derived from KDC203. Based on their chemical characteristics, it is to be predicted that each one of them might be more hydrophobic than KDC203. Naturally, as the hydrophobicity increases further, so does the insolubility of compounds in physiological environments, placing a practical limit on chemical substitutions that will need to be empirically evaluated. When assessing the potency of any of these compounds *in vivo*, it needs to be considered that wild-type mice and rats express an Atp1a1 α subunit that is far less responsive to CGs than their human ortholog. Further evaluations of the merits that KDC203 may hold for the treatment of prion diseases seem warranted.

## MATERIALS AND METHODS

### Sequence alignment and identification of residues lining CG binding pocket

Sequence identity assessments made use of available sequences for *S. scrofa* AT1A1 (P05024), *H. sapiens* ATP1A1 (P05023), ATP1A2 (P50993), and ATP1A3 (P13637) using the Basic Local Alignment Search Tool (BLAST), allowing both the filtering of low complexity regions and the introduction of gaps, and using the following settings: E-threshold: 0.001; matrix: BLOSUM62. The multiple sequence alignment algorithm Clustal O (Version 1.2.4) was used to determine homologous residues lining the CG binding pocket.

### Structural modeling and assessment of docking and BBB penetrance scores

Because no crystal structure of a human NKA is available, homology models of human α subunits were built on available structures for pig NKAs. Modeling focused on PDB entry 4HYT because it was co-crystallized with a cardenolide CG, namely ouabain, and a Mg^2+^ ion at a resolution of 3.4 Å. The availability of another porcine NKA structure, co-crystallized with a bufadienolide, namely bufalin, and two K^+^ ions at a similar resolution (4RES) was initially compared to learn about the degree to which CGs with five-versus six-membered lactone rings and different conformational states of the NKA complex affect the CG binding pose. Based on primary sequence comparisons, it was hypothesized that human NKAs comprising ATP1A3 bound to a cardenolide will exhibit the same macro 3D structure as porcine NKAs. Atomic models of human ATP1A3 bound to ouabain or oleandrin were modeled using soft minimization of ouabain geometries and amino acid residues within 5Å around it. Binding free energies were approximated using the generalized Born surface area (GBSA) method. Docking studies within the human ATP1A3 model reproduced the binding mode of ouabain reported in the literature and hypothesized a highly similar binding pose for oleandrin.

### Design of chemically accessible oleandrin derivatives

Scaffolds 1–5 depicted (**Fig 3**) were designed based on their potential semi-synthetic accessibility (1-5 step sequences) from the commercially available natural products oleandrin and gitoxin. It was envisioned that compounds depicted by Scaffold 1 would be derived from oleandrin by acyl group modification at the C16 position. The compounds denoted by Scaffolds 2 and 3 are derivatives of gitoxin that can be obtained through the introduction of the C16 acyl group followed by a deglycosylation/reglycosylation sequence. The compounds denoted by Scaffolds 4 and 5 represent derivatives of oleandrin, with Scaffold 4 compounds accessible through a known oxidation of the 4’ position [61, 62] followed by C16 acyl group adjustment. The Scaffold 5 can be derived from Scaffold 4 through a known reductive amination [46]. All compounds were assessed and ranked based on their predicted Rankovic MPO.v2 and *in silico* docking scores.

### Synthesis of shortlisted oleandrin derivative

The oleandrin derivative KDC203 was synthesized by using a well-established protocol for the PCC oxidation of oleandrin [46, 61]. Following the published purification protocols, the synthetic material was found to be >95% pure and was used as such for the subsequent biological studies.

### Characterization of KDC203 by mass spectrometry

A 780 μM KDC203 stock solution in dimethyl sulfoxide (DMSO) was stored at 37°C and sampled after 1, 3, 7 and 14 days. Each sample was frozen at collection. Upon thawing, the samples were spiked with deuterated oleandrin in DMSO then dried in a centrifugal concentrator. The remaining solids were disolved in 0.1% formic acid in a 1:1 methanol:water mix, making the concentration of deuterated oleandrin 2 μM and that of KDC203 equivalent to 10 μM.

The solutions were infused at 2 µL per min through a heated electrospray ionization source coupled to an Oribtrap Fusion mass spectrometer. Mass spectra were collected on the orbitrap mass analyzer at a nominal resolution of 60,000 with the automatic gain control target held at 4.0 e^5^, and 30 seconds of data from each sample were averaged for quantification.

### Animal husbandry

The use of *Atp1a1^s/s^* mice [52] was kindly authorized by Dr. Jerry B Lingrel (Department of Molecular Genetics, Biochemistry and Microbiology, University of Cincinnati College of Medicine, Cincinnati, Ohio 45267-0524, USA) and the animals were obtained from NOD. The mice were kept at no more than five mice per cage on a 12-hour artificial day/night cycle. Cage changes occurred once a week. The mice were subjected to daily health checks for activity and overall appearance. Drinking water was available *ad libitum* and the mice were fed a protein chow (18%). All animal procedures were in accordance with the Canadian Council on Animal Care, reviewed and authorized by the University Health Network Animal Care Committee and approved under Animal Use Protocol 6182.

### Tritium-based comparison of bioavailability of oleandrin and KDC203

Tritiated KDC203 with a specific activity of 1.6 Ci/mmol and a concentration of 1.0 mCi/mL was obtained through a customized radiolabeling request (Moravek Inc, Brea, California, USA). To minimize rapid back exchange of tritium with available protons, the radiolabeled product was overwhelmed with non-labeled water and organic solvent three times. This procedure achieved an amount of exchangeable tritium <0.1%. The certificate of analysis attested to a 100% radiochemical purity on the basis that 99.68% of radiolabeled material co-eluted with the unlabeled reference standard on a Zorbax SX 4.6 x 250 mm column (catalog number 959990-912, Agilent Technologies Canada, Mississauga, Ontario, Canada) using a mobile phase composed of 26% methanol, 26% acetonitrile, 48% water and 0.1% TFA (v/v).

Subcutaneous bolus injections of *Atp1a1^s/s^* mice with these tritiated compounds, were followed 24 hours later by tissue dissection. To this end, the mice were deeply anesthetized by isoflurane inhalation, then exsanguinated (with concomitant blood collection) by two-minute transcardiac perfusion with PBS. Brain, liver, kidney, and heart tissues were rapidly dissected. Tissues were homogenized in a RIPA buffer composed of 1% NP40, 0.5% DOC, 0.1% SDS, 150 mM NaCl, 100 mM Tris (pH 8.3) in 2 mL straight-sided test tubes with O-ring seals using 3x 1 min bead beating pulses (Beadbeater-8, Biospec, Bartlesville, Oklahoma, USA) and 200 mg of 1 mm diameter zirconia beads (catalog number 11079110zx, Biospec). Tissue extracts containing tritiated CGs were transferred to vials preloaded with scintillation fluid, and their counts were determined by scintillation counting (LS6500 Liquid Scintillation Counter, Beckman Coulter, Brea, California, USA). Concentration series, obtained by spiking known amounts of the tritiated compounds into extracts of the respective tissues from naïve mice, were used to translate experimentally obtained scintillation counts into CG concentrations. These analyses yielded total brain (C_total,brain_), plasma (C_total,plasma_), liver (C_total,liver_) and kidney (C_total,kidney_) concentrations as well as values for the total amount of the CGs of interest in the respective tissues (A_brain_).

### Bidirectional MDCK assay

The transcellular fluxes of CGs of interest were analyzed using Madin-Darby canine kidney Type II (MDCKII) cells seeded onto semi-permeable membranes. The permeability screen was conducted as a bidirectional assay encompassing the investigation of flux from the apical (A) to the basolateral (B) side and vice versa. This assay design provides the ability to elucidate the influence of extrusion pumps on the total flux. Here, the focus was on the human ATP-dependent translocase (ABCB1), often referred to by its alternative names P-glycoprotein 1 or multidrug resistance protein 1 (MDR1), which has been reported to extrude several CGs from the brain [53, 63]. We followed the convention of considering compounds MDR1 substrates if their efflux ratios, derived by dividing coefficients for B to A fluxes with those for A to B fluxes, exceed values of 2. According to complementary federal drug administration (FDA) guidelines, the addition of Zosuquidar, a highly selective, potent, and non-competitive MDR1 inhibitor [64], should decrease the efflux ratios by at least 50% to reliably classify compounds of interest as MDR1 substrates.

MDCKII cells in which the endogenous canine *MDR1* (*cMDR1*) gene was replaced with a heterologous human MDR1 (hMDR1) (catalog number MTOX1303, Millipore Sigma, Burlington, Massachusetts, USA) were maintained in high glucose Dulbecco’s Modified Eagle Medium (catalog number 11965092, Thermo Fisher Scientific, Waltham Massachusetts, USA) supplemented with 10% fetal bovine serum, 1x non-essential amino acids (catalog number 11140035, Thermo Fisher Scientific), penicillin/streptomycin (100 U/ml and 100 µg/ml, respectively; catalog number 15140122, Thermo Fisher Scientific) and 1x GlutaMAX (catalog number 35050061, Thermo Fisher Scientific). The cultures were grown at 37°C in an atmosphere of 5% CO_2_ and 95% relative humidity. The media were replaced every 48 hours, and the cells were passaged at 90% confluency using Trypsin/EDTA solution (25200072, Thermo Fisher Scientific).

12-well cell culture inserts with 0.4 µm pore size polycarbonate membranes (catalog number 140652, Thermo Fisher Scientific), deployed for the transport assay, were preincubated with cell culture media (see above) for 60 min to improve cell attachment. To monitor monolayer formation, the transepithelial electrical resistance (TEER) was measured with the EVOM2 meter (World Precision Instruments, Sarasota, Florida, USA). The measured resistance was multiplied by membrane surface area (Ω · cm^2^) and normalized by background resistance from a non-seeded insert. Five days after cell seeding, the confluent cell layers were twice washed and preincubated for 30 min at 37°C with Hank’s balanced salt solution (HBSS) (catalog number 14025, Thermo Fisher Scientific) supplemented with HEPES (catalog number 15630080, Thermo Fisher Scientific), hereafter referred to as HBSS-HEPES. The membrane integrity was further evaluated by measuring Lucifer Yellow (LY) (catalog number 07-200-156, Thermo Fisher Scientific) transepithelial leakage. To this end, inserts were transferred to a new 12-well plate containing HBSS-HEPES, and 100 µM LY solution was added to the inserts. After moderate shaking of the Nunc plates at 37°C for one hour at 70 rpm (to prevent the formation of an unstirred layer), the concentration of leaked LY was measured on the basolateral side at excitation/emission wavelengths of 428/536 nm using a SpectraMax I3 plate reader (Molecular Devices, San Jose, California, USA). Only monolayers exhibiting an apparent permeability (Papp) of Lucifer Yellow of less than 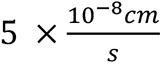 and TEER values above 97 Ωcm^2^, consistent with published resistance values [53, 65], were included in subsequent analyses. Next, inserts that passed inclusion criteria were transferred to new 12-well plates. The apical (A, insert) volume was 500 µL and the basolateral (B, well) volume was 1000 µL. For monitoring A➔B versus B➔A transport, a final concentration of 1 µM of tritiated CG of interest, dissolved in HBSS-HEPES, was added to the inserts or basolateral well, respectively. When investigating the influence of MDR1 inhibition on transport, 5 µM Zosuquidar-HCl (catalog number SML1044, Sigma-Aldrich, St. Louis, Missouri, USA) was added to both compartments. All transport experiments were conducted in biological triplicates under moderate shaking (70 rpm) for a duration of 77 minutes. To determine Papp, efflux ratio, and recovery, three aliquots were extracted from both compartments and their radioactivity counts measured with a LS6500 Liquid Scintillation Counter (Beckmann Coulter). Recovery values over 70% were recorded for all biological replicates. The following formula was used to calculate the apparent permeability coefficient Papp:

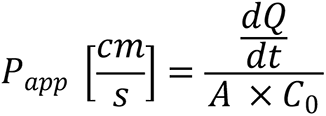

In this equation the term *dQ/dt* describes the amount of compound in the receiver compartment after a certain incubation time and therefore the rate of permeability*. A* represents the membrane surface area and *C_0_* is the starting concentration in the donor compartment.

The efflux ratio was calculated as the fraction of the Papp coefficients from the two transport types B➔A and A➔ B:

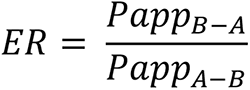

To assess the reliability of the experimental results, recovery was determined as following, where *C_f_* describes the final concentration after incubation:

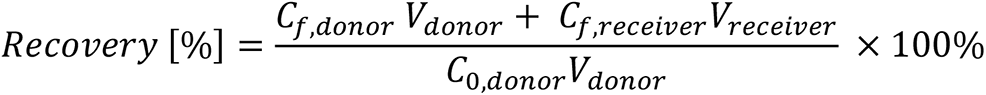

### Rapid equilibrium dialysis

Rapid equilibrium dialyses (RED) were undertaken to determine unbound fraction of oleandrin and KDC203 in brain (f_u,brain_) and plasma (f_u,plasma_). More specifically, postmortem brain tissue from *Atp1a1^s/s^* mice, guinea pigs and human frontal lobes were employed in these analyses. The origins of the human frontal lobe material were described before [66]. Briefly, this brain tissue is held in the brain bank of the Tanz Centre for Research of Neurodegenerative Diseases that was adopted from a former Canadian Brain Tissue Bank at the Toronto Western Hospital. The specific frontal lobe tissue used for these analyses originated from one male and one female who had died in their 70s of non-dementia causes. All tissues were homogenized in PBS and spiked with the respective tritiated CGs before being equilibrated against PBS buffer in a RED device (catalog number 89809, Thermo Fisher Scientific). Plasma was collected from *Atp1a1^s/s^* mice and guinea pigs. The RED device was placed in a shaker orbiting at 225 rpm for five hours. Samples were then transferred from the RED device into scintillation vials for scintillation counting. The *in vitro* data from these RED analyses revealed the diluted unbound fraction (f_u,d_) of the respective CGs by determining the concentration of the tritiated CGs through scintillation counting of volume aliquots harvested from the red-colored sample input side and white-colored sample output side. Since plasma samples were undiluted, f_u,d,plasma_ equals f_u,plasma_ [67]:

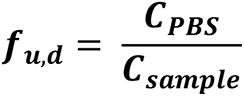

To account for the dilution of the brain homogenates, the following equation corrected for the dilution factor D [68]:

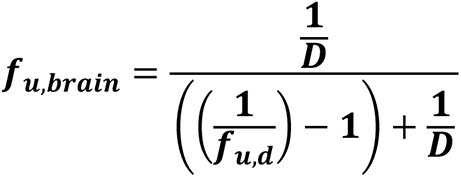

The RED-based measurements of the unbound fractions (f_u_) of CGs in brain and plasma homogenates was used to estimate the unbound CG concentrations in the respective organs (C*_u_*) that was achieved following the subcutaneous bolus injections of tritiated CGs into *Atp1a1^s/s^*mice (see above):

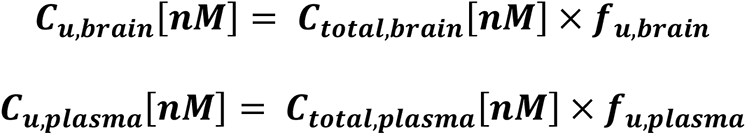

With this information in hand, the unbound drug partitioning coefficient *K_p,uu_* was computed from the fraction of unbound CGs in brain and plasma [59]:

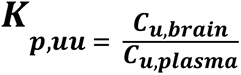

Finally, the unbound volume of distribution in the brain (V_u,brain_) was calculated by dividing the total amount of compound in the brain (A*_brain_*) and the unbound fraction in the brain (*C_u,brain_*):

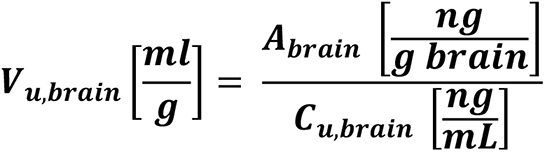

### Treatment of ReN VM cells with KDC203

ReN VM Human Neural Progenitor Cells (catalog number SCC008, Millipore Sigma) were grown in their undifferentiated form on 20ug/mL Cultrex reduced growth factor basement membrane (catalog number 3433, R&D Systems, Minneapolis, Minnesota, USA) in DMEM/F12 (catalog number 11320033, Thermo Fisher Scientific) based media with 0.22 micron filter sterilized 20 ng/mL human basic fibroblast growth factor (bFgF)(catalog number RKP09038, Reprokine, St Petersburg, Florida, USA), 20ng/mL human recombinant epidermal growth factor (EGF)(catalog number RKP01133, Reprokine), 10 units/mL heparin Na^+^ salt (catalog number 3149-10KU, Sigma-Aldrich), 2% (v/v) B-27 supplement (catalog number 17504044, Gibco / Thermo Fisher Scientific), 1% (v/v) Glutamax (catalog number 35050061, Gibco / Thermo Fisher Scientific) and 1% (v/v) non-essential amino acids (catalog number 11140050, Gibco / Thermo Fisher Scientific). Differentiation into a co-culture of non-dividing cells that exhibit neuronal and astrocytic characteristics was initiated with the removal of growth factors and heparin salt from media. Cells were differentiated for 7 days with replacement of media every 2 days. Treatment was initiated on day 8 of differentiation with the addition of nanomolar media concentrations of KDC203 or oleandrin (both solubilized in DMSO) to the differentiation media. Treatment continued for 7 days during which 50% of media containing the appropriate concentration of the drug were replenished daily. Vehicle-treated cells received the same concentration of DMSO as CG-treated cells.

### Antibodies

Immunoblotting analyses of PrP relied on the monoclonal 3F4 antibody (catalog number MAB1562, Millipore Sigma) which recognizes the 109-112 amino acid epitope of human PrP. Antibodies against the NKA subunits consisted of anti-ATP1A1 (catalog number ab7671, Abcam Inc, Toronto, Ontario, Canada), anti-ATP1A2 (catalog number ab9094-I, Abcam Inc), anti-ATP1A3 (catalog number MA3-915, Thermo Fisher Scientific) and anti-ATP1B1 (catalog number GTX113390, GeneTex, Irvine, California, USA). Neuronal and astrocytic markers were detected with the anti-NeuN antibody (catalog number EPR12763, Abcam Inc), the anti-Tuj-1 antibody (catalog number ab78078, Abcam Inc) and the anti-GFAP monoclonal (131-17719) antibody (catalog number A-21282, Thermo Fisher Scientific).

### Western blot analyses

Cells were lysed in ice-cold buffer containing 1% NP40, 150 mM Tris-HCL (pH 8.3) and 150 mM NaCl supplemented with a protease inhibitor cocktail (catalog number 11836170001, Sigma-Aldrich) and phosSTOP phosphatase inhibitor cocktail (catalog number 4906845001, Millipore Sigma). Insoluble cellular debris was removed following a 5-minute slow speed centrifugation step (2000 g), followed by a 15-minute fast speed centrifugation (16000 g). Protein concentrations of the supernatants were determined via the bicinchoninic acid assay (BCA) using the Pierce BCA Protein Assay Kit (catalog number 23225, Thermo Fisher Scientific). Following BCA-based protein concentration measurements, protein levels were adjusted to identical concentrations with cell lysis buffer. Samples for immunoblot analyses were denatured and reduced in 1x Bolt LDS sample buffer (catalog number B0007, Thermo Fisher Scientific) and 2.5% β-mercaptoethanol (catalog number M6250, Sigma-Aldrich), then heated at 95°C for 10 minutes and briefly cooled on ice, immediately prior to gel loading. Proteins were separated by SDS-PAGE on 10% Bolt Bis-Tris gels (catalog number NW00105BOX, Thermo Fisher Scientific), then transferred to 0.45 micron PVDF membranes (catalog number IPVH00010, Millipore Sigma), and unspecific binding was reduced by blocking in 5% skimmed milk (catalog number SKI400, BioShop, Burlington, Ontario, Canada) for 1 hour at room temperature. Membranes were then incubated in primary antibodies diluted in 5% skimmed milk and left overnight at 4°C with gentle rocking. The following day, membranes were washed three times in 1x Tris-buffered saline with 0.1% Tween20 (catalog number TWN508, BioShop) (TBST), then incubated with the corresponding horseradish peroxidase (HRP)-conjugated secondary antibodies, diluted in 5% skimmed milk, and left on the rocker for 1 hour at room temperature. Membranes were again washed three times in 1x TBST, then incubated with enhanced chemiluminescent (ECL) reagent (catalog number GERPN2232, GE HealthCare) for 1 minute. Membranes were exposed to autoradiography film (catalog number CLMS810, MedStore, Toronto, Ontario, Canada) and developed via a film developer.

### Viability, intracellular Ca^2+^, and ATP content in cells exposed to KDC203 versus oleandrin

We used three assays for comparing the cellular health and metabolism of ReN VM cells, treated with KDC203 versus oleandrin. For each of these assays, the cells were passaged to clear-bottom 96-well tissue culture plates and differentiated for one week by the withdrawal of growth factors as described above.

To initiate a cell viability assay, the cells were washed with PBS (catalog number D8537-500 ML, Sigma-Aldrich), then incubated for 20 minutes with 1 µM calcein-AM (catalog number 17783, Sigma-Aldrich) at 37°C, followed by incubation in PBS supplemented with bovine serum albumin (BSA) (catalog number ALB001.50, BioShop) at 3% (w/v) to fully de-esterify AM esters. The subsequent analysis occurred in a microplate reader at excitation/emission wavelengths of 486 nm and 516 nm.

To measure cellular ATP levels, CellTiter-Glo (catalog number G7570, Promega, Madison, Wisconsin, USA) was 1:4 diluted in PBS and added at a 1:1 (v/v) ratio to the growth factor-depleted cell differentiation medium covering the cells. Following a 2-minute agitation at room temperature that caused the cells to lyse, the data from 1-second luminescence recordings were integrated and taken to reflect cellular ATP content.

The intracellular Ca^2+^ concentration was assayed for 20 min at 37°C with 2 µM Fluo-4-AM (catalog number F14201, Thermo Fisher Scientific) in PBS, supplemented with 3% (w/v) BSA. Following a PBS wash and an additional 30-minute incubation in BSA-supplemented PBS to fully de-esterify AM esters, the excitation and emission were recorded at 486 and 516 nm wavelengths in a microplate reader, respectively, and taken to reflect the intracellular Ca^2+^ content.

The oleandrin measurements shown here were already included in a previous report (along with results for digoxin, not shown here) [18] and were at the time generated side-by-side with the data for KDC203. In assembling this report, we chose to again include the oleandrin results, this time next to the KDC203 data, to facilitate their direct comparison.

### Statistical analyses

Comparisons of steady-state protein levels by western blot analyses made use of three biological replicates. Six biological replicates for each CG concentration tested were used to determine the cellular health and metabolism in cells using the calcein-AM, CellTiter-Glo, and Fluo-4-AM assays. Biological triplicates were deployed for the statistical analyses of pharmacokinetic data comparing KDC203 and oleandrin in brain homogenates from distinct species. The two-tailed ^t^-test was employed to the statistical analyses of all cohorts based on the assumption of unknown variances of grouped samples. Following general convention in biomedical research, one, two, and three asterisks indicate significant p-values of <0.05, <0.005, and <0.0005, respectively. The abbreviation ‘ns’ is shown when p-values failed to meet the significance threshold.

### Ethics statement

The use of human ReN VM and T98G cell lines for experiments shown in this report was authorized by the office for Environmental Health a Safety at the University of Toronto, Toronto, Ontario, Canada (Biosafety Permit 208-S06-2). The mouse work was approved by the Animal Care Committees of the University of Toronto and the University Hospital Network (Animal Use Protocol 6182).

## ACKNOWLEDGMENTS

The authors acknowledge the generous sharing of *Atp1a1^s/s^* mice by the late Dr. Jerry B Lingrel and are most grateful to the Krembil Foundation, the Borden Rosiak family, and the Arnold Irwin family for their outstanding support of this project. PN acknowledges support by NIH R35 GM136341.

## AUTHOR CONTRIBUTIONS

Conceived and designed the experiments: SE, TZ, XW, DW, CS, SM, PN, GS. Performed the experiments: SE, TZ, DW, XW, CS, SM. Analyzed the data: SE, TZ, XW, DW, CS, SM, PN, GS. Contributed reagents/materials/analysis tools: ND, JBL, GS. Wrote the first version of paper: GS. Edited the manuscript: All authors.

## SUPPLEMENTAL MATERIAL

**S1 Fig. MS-based evidence of successful synthesis and purity of novel lead compound**

(A) Design of KDC203 stability test undertaken at 36.6°C (corresponding to the normal body temperature of mice). (B) Exemplary mass spectrum of KDC203. (C) Isotopic envelop of KDC203 with a freshly dissolved internal standard CG spiked into the sample following extended incubation but prior to sample processing for LC-MS. KDC203 continued to give rise to the base peak of the mass spectrum and no detectable degradation relative to the internal standard was detectable during the 14-day incubation. (D) Table summarizing intensities and ratios of intensities of monoisotopic peaks for KDC203 and the internal standard CG at the selected time intervals.

**S2 Fig. Crude estimation of human cerebrum levels achievable with KDC203**

(A) Brain and plasma levels of oleandrin achieved in wild-type mice 24 hours following intraperitoneal injection of a one-time dose of 3 µg/g [31]. (B) Cerebrum and plasma levels of oleandrin observed in cats following once daily intravenous injection of 3 µg/kg [30]. (C) Average total brain concentration of oleandrin achieved in *Atp1a1^S/S^* mice 24 hours following subcutaneous injection of a one-time dose of 0.6 µg/g (this study). (D) Plasma levels of oleandrin observed following daily oral administration for a duration of 8 days of 4.4 µg/kg of oleandrin present in a *Nerium oleander* extract [70]. (E) Average total brain concentration of KDC203 achieved in *Atp1a^S/S^* mice 24 hours following subcutaneous injection of a one-time dose of 0.6 µg/g (this study). When estimating unbound KDC203 levels achievable in human brains, we can draw on three data points that seem most pertinent: 1. Cerebrum levels of oleandrin in the brain of cats were 6-fold higher than their plasma levels after daily administration. If a similar daily administration regime reached 3 nM oleandrin levels in the human plasma, then this might indicate that brain levels could have been as high as 18 nM. 2. Our direct comparison of oleandrin and KDC203 brain levels in mice suggests that KDC203 might reach 1.5-fold higher total brain concentrations. If this relative propensity translates to humans, then this indicates that 18 x1.5 = 27 nM total brain concentrations of KDC203 might be attainable in human brains following an oral daily dosing regimen. 3. The RED analyses undertaken in this study with human brain samples, revealed that a ratio of 0.395 of the total KDC203 brain concentration might be available in unbound form for engagement with NKAs.

