## Supplemental Material for "Identification of a cardiac glycoside exhibiting favorable brain bioavailability and potency for reducing levels of the cellular prion protein"

### S1 Fig

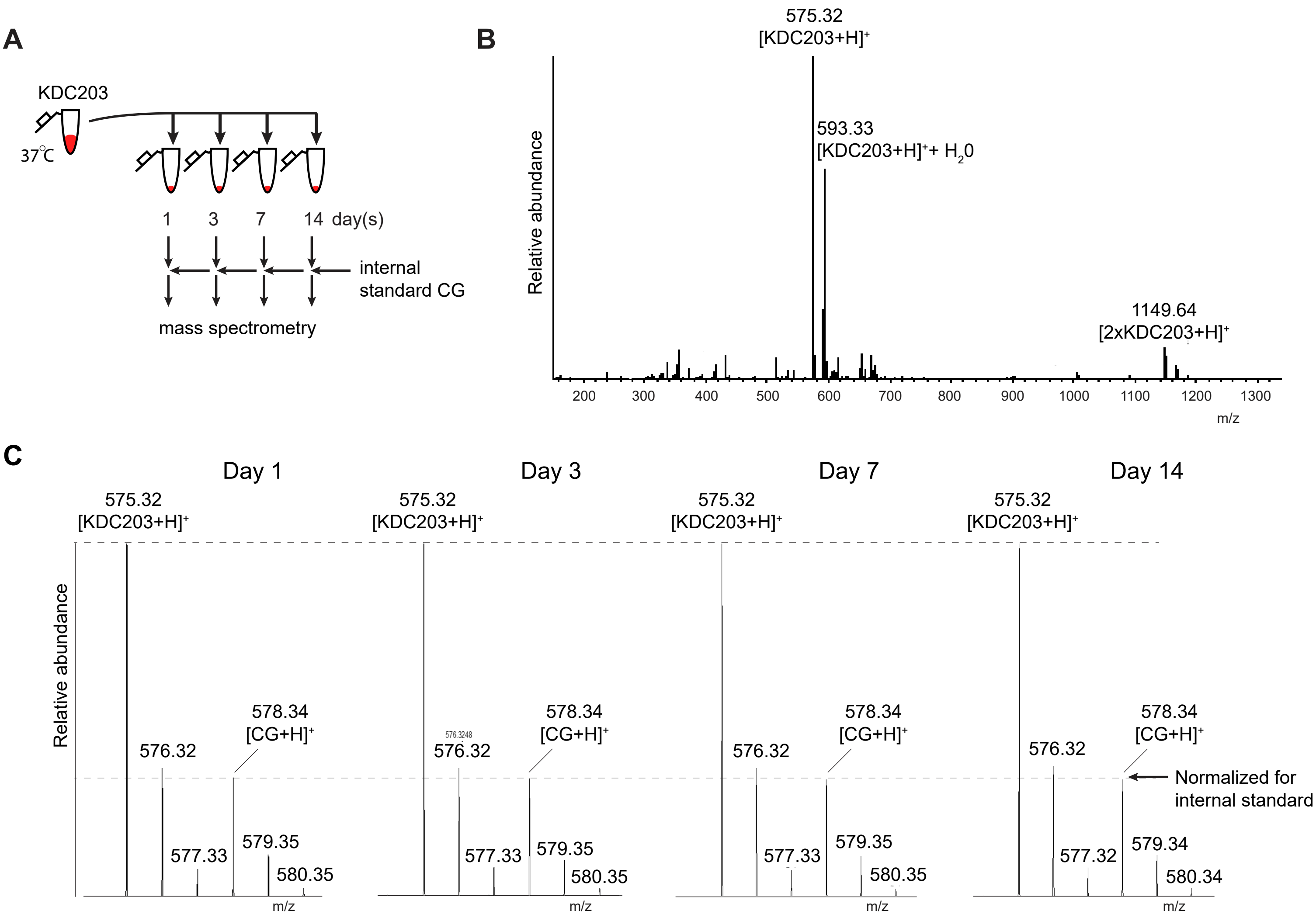


**D**

| Days at 37°C | Intensity of monoisotopic internal standard [CG+H]$^+$ at 578.34 Th (counts) | Intensity of monoisotopic lead compound [KDC203+H]$^+$ at 575.32 Th (counts) | Intensity ratio of monoisotopic lead compound [KDC203+H]$^+$ and internal standard [CG+H]$^+$ | Normalized intensity ratio of monoisotopic lead compound [KDC203+H]$^+$ and internal standard [CG+H]$^+$ |
|---|---|---|---|---|
| 1 | 3.20E+06 | 9.67E+06 | 3.02 | 1.00 |
| 3 | 3.03E+06 | 1.01E+07 | 3.33 | 1.10 |
| 7 | 5.40E+07 | 1.66E+08 | 3.07 | 1.02 |
| 14 | 3.22E+07 | 1.07E+08 | 3.32 | 1.10 |

### S2 Fig

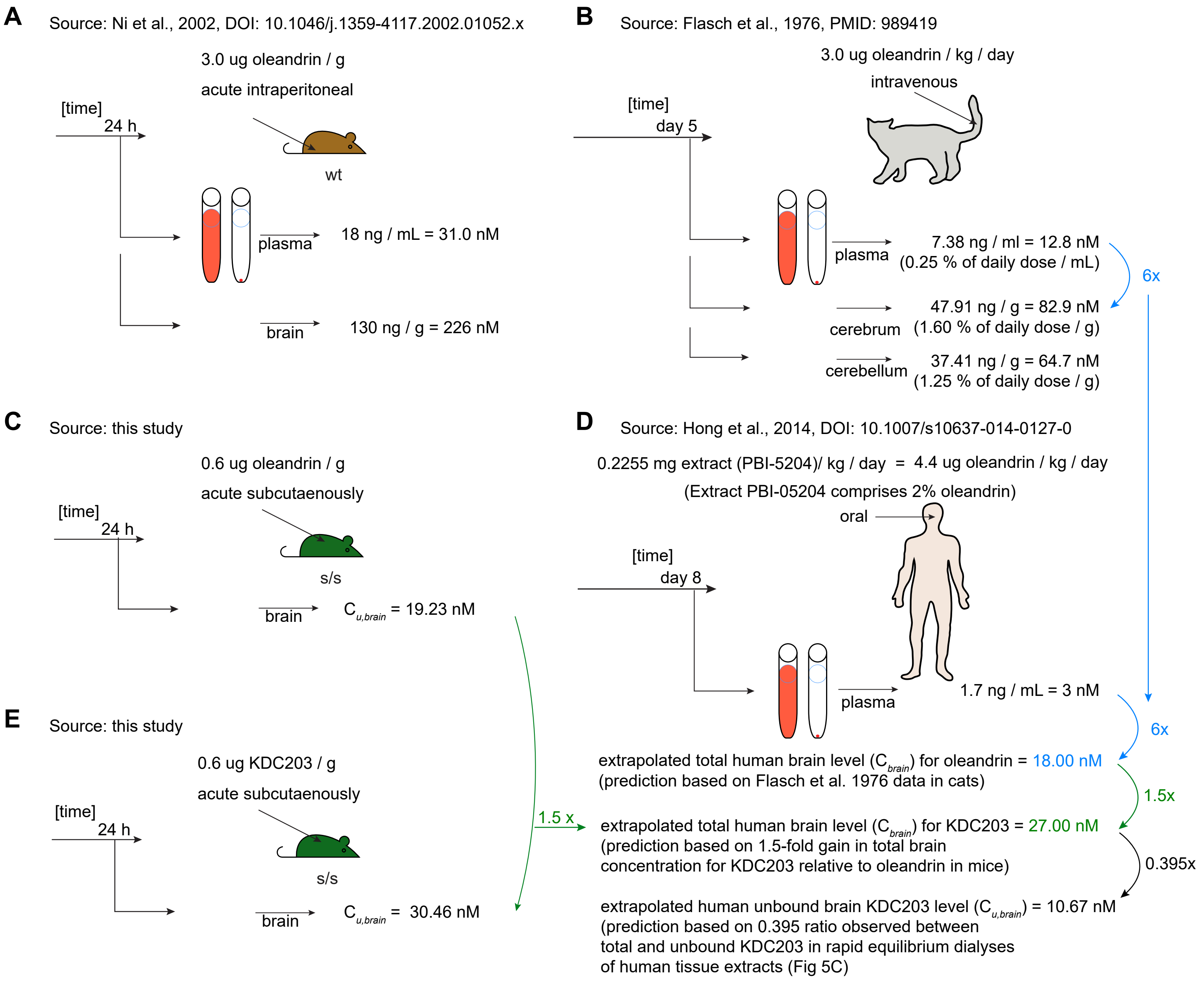

A
Source: Ni et al., 2002, DOI: 10.1046/j.1359-4117.2002.01052.x
3.0 ug oleandrin / g
acute intraperitoneal
[time]
24 h
wt
plasma
18 ng / mL = 31.0 nM
brain
130 ng / g = 226 nM
B
Source: Flasch et al., 1976, PMID: 989419
3.0 ug oleandrin / kg / day
intravenous
[time]
day 5
plasma
7.38 ng / ml = 12.8 nM
(0.25 % of daily dose / mL)
6x
cerebrum
47.91 ng / g = 82.9 nM
(1.60 % of daily dose / g)
cerebellum
37.41 ng / g = 64.7 nM
(1.25 % of daily dose / g)
C
Source: this study
0.6 ug oleandrin / g
acute subcutaenously
[time]
24 h
s/s
brain
C_{u,brain} = 19.23 nM
D
Source: Hong et al., 2014, DOI: 10.1007/s10637-014-0127-0
0.2255 mg extract (PBI-5204)/ kg / day = 4.4 ug oleandrin / kg / day
(Extract PBI-05204 comprises 2% oleandrin)
oral
[time]
day 8
plasma
1.7 ng / mL = 3 nM
6x
extrapolated total human brain level (C_{brain}) for oleandrin = 18.00 nM
(prediction based on Flasch et al. 1976 data in cats)
1.5x
1.5 x
extrapolated total human brain level (C_{brain}) for KDC203 = 27.00 nM
(prediction based on 1.5-fold gain in total brain
concentration for KDC203 relative to oleandrin in mice)
0.395x
extrapolated human unbound brain KDC203 level (C_{u,brain}) = 10.67 nM
(prediction based on 0.395 ratio observed between
total and unbound KDC203 in rapid equilibrium dialyses
of human tissue extracts (Fig 5C)
E
Source: this study
0.6 ug KDC203 / g
acute subcutaenously
[time]
24 h
s/s
brain
C_{u,brain} = 30.46 nM
